# Single-genome analysis reveals heterogeneous association of the Herpes Simplex Virus genome with H3K27me2 and the reader PHF20L1 following infection of human fibroblasts

**DOI:** 10.1101/2023.12.03.569766

**Authors:** Alison K Francois, Ali Rohani, Matt Loftus, Sara Dochnal, Joel Hrit, Steven McFarlane, Abigail Whitford, Anna Lewis, Patryk Krakowiak, Chris Boutell, Scott B. Rothbart, David Kashatus, Anna R Cliffe

## Abstract

The fate of herpesvirus genomes following entry into different cell types is thought to regulate the outcome of infection. For the Herpes simplex virus 1 (HSV-1), latent infection of neurons is characterized by association with repressive heterochromatin marked with Polycomb silencing-associated lysine 27 methylation on histone H3 (H3K27me). However, whether H3K27 methylation plays a role in repressing lytic gene expression in non-neuronal cells is unclear. To address this gap in knowledge, and with consideration that the fate of the viral genome and outcome of HSV-1 infection could be heterogeneous, we developed an assay to quantify the abundance of histone modifications within single viral genome foci of infected fibroblasts. Using this approach, combined with bulk epigenetic techniques, we were unable to detect any role for H3K27me3 during HSV-1 lytic infection of fibroblasts. In contrast, we could detect the lesser studied H3K27me2 on a subpopulation of viral genomes, which was consistent with a role for H3K27 demethylases in promoting lytic gene expression. This was consistent with a role for H3K27 demethylases in promoting lytic gene expression. In addition, viral genomes co-localized with the H3K27me2 reader protein PHF20L1, and this association was enhanced by inhibition of the H3K27 demethylases UTX and JMJD3. Notably, targeting of H3K27me2 to viral genomes was enhanced following infection with a transcriptionally defective virus in the absence of Promyelocytic leukemia nuclear bodies. Collectively, these studies implicate a role for H3K27me2 in fibroblast-associated HSV genome silencing in a manner dependent on genome sub-nuclear localization and transcriptional activity.

**Importance:** Investigating the potential mechanisms of gene silencing for DNA viruses in different cell types is important to understand the differential outcomes of infection, particularly for viruses like herpesviruses that can undergo distinct types of infection in different cell types. In addition, investigating chromatin association with viral genomes informs on the mechanisms of epigenetic regulation of DNA processes. However, there is growing appreciation for heterogeneity in the outcome of infection at the single cell, and even single viral genome, level. Here we describe a novel assay for quantifying viral genome foci with chromatin proteins and show that a portion of genomes are targeted for silencing by H3K27me2 and associate with the reader protein PHF20L1. This study raises important questions regarding the mechanism of H3K27me2-specific targeting to viral genomes, the contribution of epigenetic heterogeneity to herpesvirus infection, and the role of PHF20L1 in regulating the outcome of DNA virus infection.

## Introduction

The genomes of DNA viruses, especially those that replicate in the nucleus, have an intimate association with host chromatin. Herpesviruses are double-stranded DNA viruses that can undergo both lytic/productive replication and establish long-term latent infections. There is growing evidence that the regulation of herpesvirus latent versus lytic infection results from the deposition of cell type-specific types of chromatin, as active euchromatin is enriched during productive *de novo* infection and reactivation, whereas repressive heterochromatin is enriched on viral genome during latent infection (1–14). However, the mechanisms that regulate the deposition of heterochromatin, and functional outcomes in different cell types for many of the herpesviruses, remain unknown.

Polycomb-mediated silencing is a type of facultative heterochromatin, characterized in large part by the enrichment of lysine 27 tri-methylation on histone H3 (H3K27me3) and is associated with multiple latent herpesviruses (1, 2, 5, 9, 10, 12, 13, 15–20). Facultative heterochromatin is more readily converted to transcriptionally active euchromatin than the more stable constitutive heterochromatin (21). Polycomb silencing is primarily established on the host genome during pluripotency and remodeled during cell specification. Hence, most data on Polycomb silencing comes from studies investigating gene silencing in stem cells and during the early stages of development (19, 22). Many herpesviruses infect differentiated host cells and Polycomb silencing of latent herpesvirus genomes is believed to promote and/or maintain repression of the viral lytic phase genes during latent infection (19). The mechanisms of Polycomb silencing may differ between pluripotent and terminally differentiated cells. Therefore, investigating the *de novo* deposition of chromatin onto incoming herpesvirus genomes has the potential to inform on the mechanisms of heterochromatin formation in different cell types.

Herpes simplex virus type I (HSV-1) is the prototype alphaherpesvirus, and infection of neurons can result in a life-long latent infection in which lytic genes are repressed. In contrast, infection of epithelial cells or fibroblasts results in productive (lytic) replication. The molecular mechanisms that regulate entry into lytic replication or silent latent infection are important to understand because HSV-1 (and the related HSV-2) latent infection causes significant morbidity and mortality. Periodically, the virus reactivates from latent infection to result in infectious virus production, which can lead to lesions at the body surface, keratitis and encephalitis. In addition to these outcomes, HSV-1 infection has been linked to the progression of late-onset neurodegenerative disease (23–27).

When HSV-1 genomes initially enter the nucleus, they are devoid of chromatin. There is evidence for rapid association of histones with incoming genomes mediated by histone chaperone proteins (28–32). During a latent infection of neurons, HSV-1 lytic genes are associated with cellular histones carrying H3K27me3 as well as di- and tri-methylation of lysine 9 on histone H3 (H3K9me2/me3) (12, 15, 19, 33–35). One model for heterochromatin association with the HSV-1 genome involves immediate recognition of incoming viral DNA by the host cell, resulting in heterochromatin-mediated silencing, which the virus must overcome for lytic replication (6, 36–40). In contrast, the virus is thought to be unable to overcome gene silencing in neurons, and latency is established (6). However, for Polycomb silencing, evidence supporting the initial deposition of H3K27me3 during lytic infection is limited. Previously, ubiquitously transcribed tetratricopeptide repeat, X chromosome (UTX) protein was shown to contribute to HSV-1 gene expression in U2OS cells (41). UTX (also known as Lysine-specific Demethylase 6A; KDM6A) is one of two H3K27-specific demethylases, the second being Jumonji domain-containing protein-3 (JMJD3 or KDM6B). A further study described transient formation of H3K9me3 and H3K27me3 on a lytic promoter immediately upon infection, followed by distinct waves of removal (36). However, the absolute levels of H3K27me3 enrichment observed in this study were orders of magnitude lower than the enrichment on host chromatin. In addition, fully determining presence of histone modifications can be problematic due to potential background enrichment in these assays, and interpretation is further complicated by appreciated issues with histone PTM antibody specificity (42). Another confounding factor is potential heterogeneity in the outcome of HSV infection, even in mitotic fibroblast cell lines (43–48). These studies suggest that sub-populations of viral genomes may associate with certain types of heterochromatin, which would be difficult to detect in bulk assays like chromatin immunoprecipitation (ChIP). Assays that can measure the enrichment for certain types of chromatin at the single genome level would therefore be beneficial to take into account this potential heteroeneity.

Previous studies have identified the presence of the constitutive heterochromatin marks H3K9me2/me3 on the viral genome during lytic infection and defined their role in gene silencing (36, 38, 49, 50). Notably, H3K9me2/me3-marked histones associate with the histone chaperone protein DAXX prior to loading onto chromatin, facilitating rapid association of repressive PTMs with viral DNA (51). DAXX also associates with Promyelocytic leukemia (PML) nuclear bodies (NBs) (also known as ND10), and previous literature supports a model in which PML-NBs are involved in constitutive heterochromatin formation (32, 52–57). Upon infection of non-neuronal cells, HSV-1 genomes rapidly associate with PML-NBs and there is evidence for silencing through H3K9me2/me3 at these bodies (32, 48). Consistent with this observation, PML-NBs repress HSV-1 lytic replication, and the viral protein ICP0 degrades PML to overcome this repression (58–64). However, histones pre-modified with methylated lysine H3K27 have not been previously detected (65). Given the association of viral genomes with PML-NBs and Daxx, and how loss of PML shifts the balance of methylation from H3K9 to H3K27 (54), it is not clear whether incoming lytic genomes could be targeted for Polycomb silencing simultaneously with H3K9 methylation. Notably, we recently reported that primary neurons are devoid of PML-NBs (66), and hence if these bodies do protect against Polycomb silencing, viral genomes would still be targeted for H3K27me3 deposition following neuronal infection. However, the presence of PML-NBs in non-neuronal cells could shift the balance of heterochromatin away from Polycomb silencing and towards initial H3K9me2/3 deposition.

The previously observed kinetics of H3K27me3 formation (approximately 36 hours), at least in pluripotent cells, also do not support a role for this modification in restricting lytic HSV-1 infection (67, 68). However, it cannot be ruled out that there are different kinetics for *de novo* H3K27me3 formation on incoming viral genomes in more differentiated cell types. Importantly, the deposition of H3K27me2 is much more rapid (69). H3K27me2 is also associated with gene silencing, and a recent study identified PHF20L1 as a repressive reader protein of this PTM (70). H3K27me2 is the most abundant H3K27 methylation state in embryonic stem cells (69) and yet the H3K27me2 association with herpesvirus genomes, and any contribution to gene silencing, have not previously been reported.

Here, we set out to define the contribution of Polycomb silencing of HSV-1 genomes to lytic gene repression in non-neuronal cells. Using a combination of epigenetic, imaging and gene expression-based approaches, we determined that HSV-1 infection in primary human foreskin fibroblasts (HFFs) does not result in Polycomb-mediated H3K27me3 deposition or its subsequent removal. Importantly, we developed a novel assay quantifying co-localization between viral genome foci and histone PTMs in the nucleus, therefore accounting for heterogeneity between different genome epigenetic structures. Combining this analysis with the inhibition of demethylases UTX and JMJD3, we found that a sub-population of genomes was associated with H3K27me2 and PHF20L1. Further, association with H3K27me2 increased on genomes of virus lacking the transactivating function of VP16 in the absence of PML-NBs, suggesting that association with PML-NBs limits H3K27me2 deposition on transcriptionally inactive genomes and instead promotes more constitutive heterochromatin formation.

## Results

### H3K27me3 does not associate with the HSV-1 genome following infection of human fibroblasts

To determine the potential contribution of H3K27 methylation mediated by Polycomb repressive complexes during HSV-1 lytic replication in non-neuronal cells, we set out to investigate the deposition of H3K27me3 on the viral genome following infection of human fibroblasts. In a previous study, data from ChIP followed by qPCR suggested a low level of H3K27me3 was present at 1-2 hours post-infection (hpi) and was later reduced, co-incident with reduced total histone association and following ICP0 expression (36). However, without comparing H3K27me3 on the viral genome to regions of host chromatin that are enriched for this PTM, it cannot be concluded whether this modification is enriched on the viral genome. In addition, since this study was published, data have emerged that many antibodies against H3K27 methylation show non-specific binding to other histone PTMs (42), and it is therefore possible that antibody cross-reactivity with other PTMs complicated interpretation of these data.

To resolve these issues, we infected HFFs with HSV-1 strain 17 Syn+ at an MOI of 3 plaque forming units (PFU) per cell and carried out CUT&RUN 2 and 4 hpi; time points chosen based on published observations that both H3K27me3 and total histone levels peak at these times and are removed by 4 hpi (31, 36). The H3K27me3 antibody used was chosen for its high target specificity, as determined by histone peptide array analysis (42) (Figure S1A). To quantify enrichment at specific promoter sequences, we analyzed the enrichment compared to the non-specific IgG control aligning to the KOS reference strain (71). CUT&RUN data for both 2 and 4 hpi show very little enrichment for H3K27me3 on viral lytic promoters, shown as the geometric means of 2 replicates (linear enrichment over IgG) scaled to the host positive control (Myt1) promoter in Figure 1. All IE and early gene promoters are shown, with a selection of late genes. In the presence of JMJD3 and UTX inhibition (GSK-J4), the host genome showed higher enrichment for H3K27me3 at 4 hpi the Myt1 promoter, in addition to an increase compared to the 2h time-point, which is consistent with previous reports (72). GSK-J4 activity was validated for uninfected HFF chromatin by western blot (Figure S2A), indicating that the lack of H3K27me3 accumulation on viral promoters is not due to a lack of inhibitor activity. In stark contrast to the host promoters, no viral lytic promoters showed notable enrichment for H3K27me3. Therefore, in bulk cultures, we were unable to detect positive enrichment for H3K27me3 on the regions of the viral genome examined.

**Figure 1.**
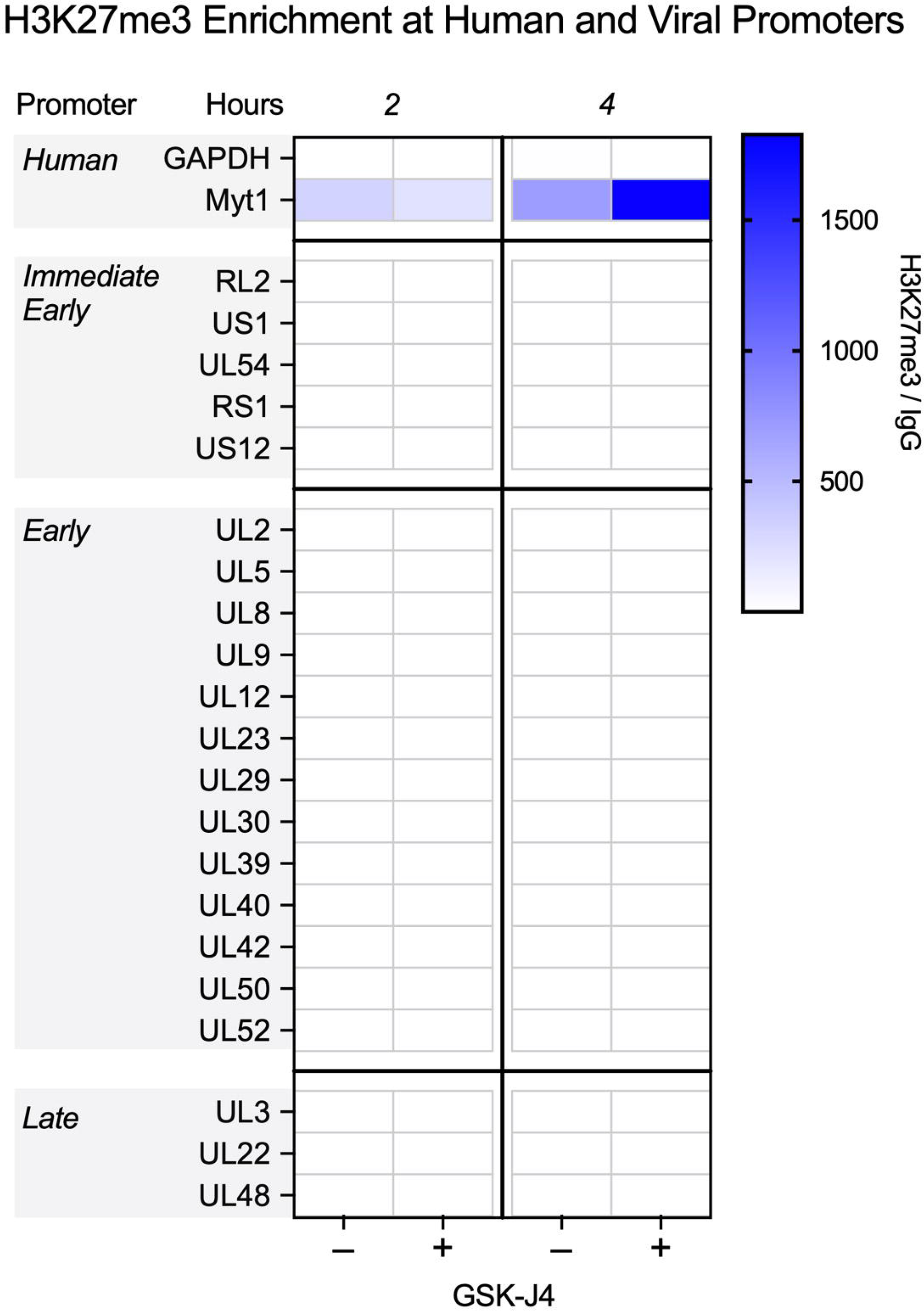
CUT&RUN during early infection shows little H3K27me3 enrichment on lytic HSV-1 chromatin. HFFs were infected at MOI 3 PFU/cell untreated or treated with 10µM GSK-J4. Cells were processed for CUT&RUN, and fragments sequenced and aligned to both human and viral genomes. The sum of coverage at defined promoter regions was used to calculate fold enrichment of H3K27me3 over IgG. The geometric mean of the fold enrichment is plotted as a heat map scaled to host gene *Myt1* (N=2).

### Analysis of individual HSV-1 genome foci for co-localization with H3K27me3 using NucSpotA

Although the CUT&RUN data suggest there was very little H3K27me3 enriched on the lytic HSV-1 genome following infection of fibroblasts, we could not rule out the possibility that a subpopulation of viral genomes associates with H3K7me3 upon fibroblast infection, which may not be detected by these bulk population-level methods. Therefore, we developed an assay that would permit the quantification of histone modifications associated with HSV-1 DNA at an individual genome, or genome spot, resolution. Importantly, other studies have observed heterogeneity in the ability of cells to support lytic replication (43, 48). Therefore, it was possible that there is heterogeneity in heterochromatin association with viral genomes following infection of fibroblasts.

We prepared viral stocks that contained EdC-labelled genomes as previously described (59). HFFs were infected with EdC-labelled HSV-1 (HSV^EdC^), and Click chemistry-based fluorescent staining (to visualize viral DNA) was carried out alongside immunostaining with the chosen histone antibody (60). To accurately quantify the enrichment of each histone PTM with the viral genome in an unbiased and high-throughput manner, we developed a custom program (NucSpotA) that measures the intensity of signal at a viral genome compared to the intensity of positive signal throughout the nucleus (Figure 2). We first validated NucSpotA by quantifying the co-localization with proteins that have been found enriched at sites of viral genomes; RNA polymerase II (Figure 3A, top) and total histone H3 (Figure 3A, middle). H3 is known to be rapidly deposited on the lytic HSV-1 genome (6, 30, 73–75), while RNA polymerase II is essential for viral lytic gene expression (76) and has been shown previously to co-localize with viral genomes (77). A higher intensity ratio represents a higher enrichment of a protein at viral genomes.

**Figure 2.**
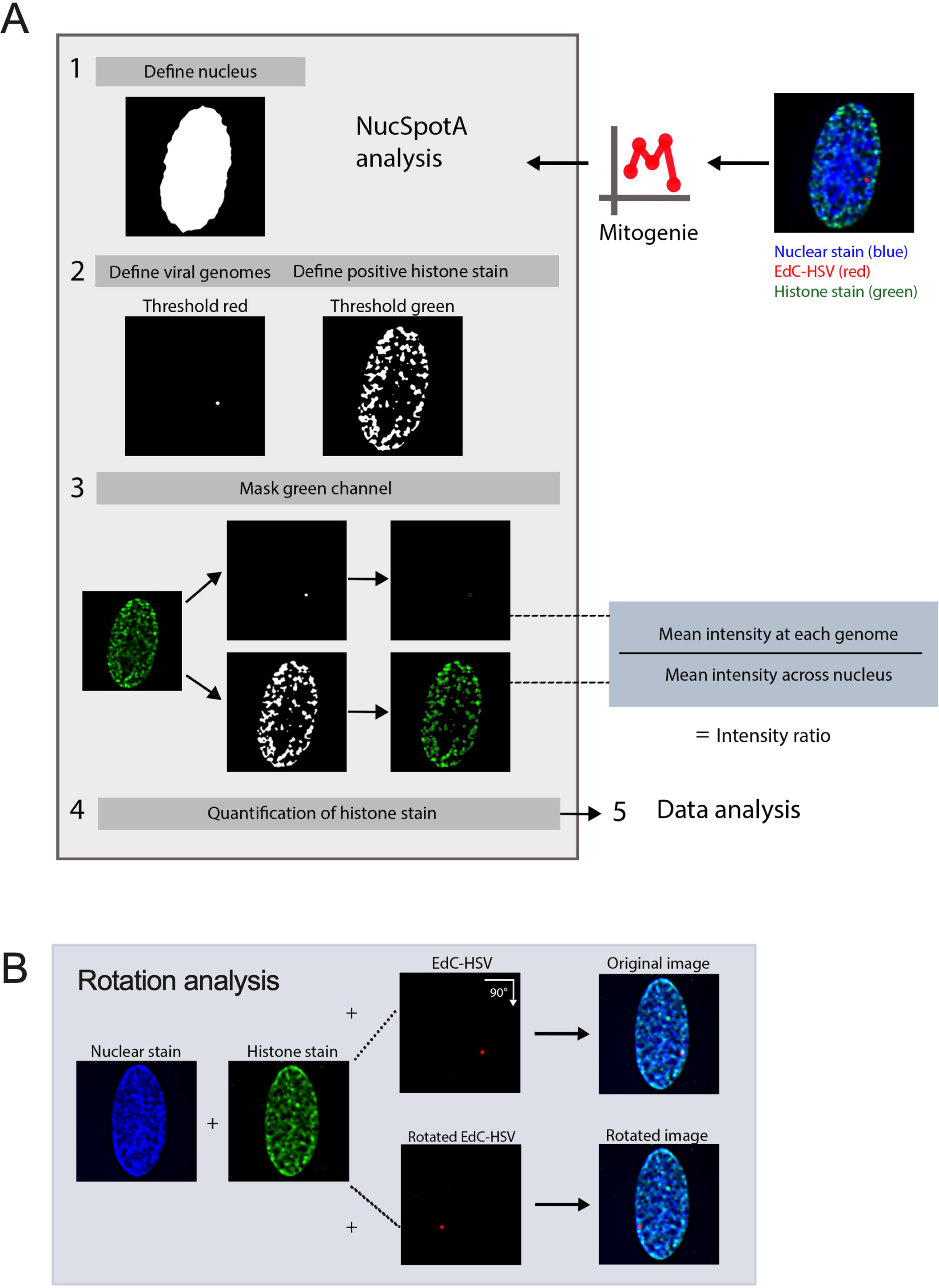
Image analysis using NucSpotA to quantify co-localization between viral genomes and an immunostained target. (A) The workflow used for a batch of images of individual infected nuclei using NucSpotA. An RGB image is thresholded to define the nucleus (blue channel), then viral genomes (red channel) and positive immunostain signal (green channel) within the nucleus. Representative images show a histone immunostain (H3K27me3). Mean intensity of the immunostain at each viral genome and across the nucleus is measured, and used to calculate an intensity ratio. (B) Rotation of the red channel relative to blue and green channels is used to generate pairs of original and rotated images. Rotation functions as random placement of viral genomes within the nucleus. Image pairs are processed in parallel.

**Figure 3.**
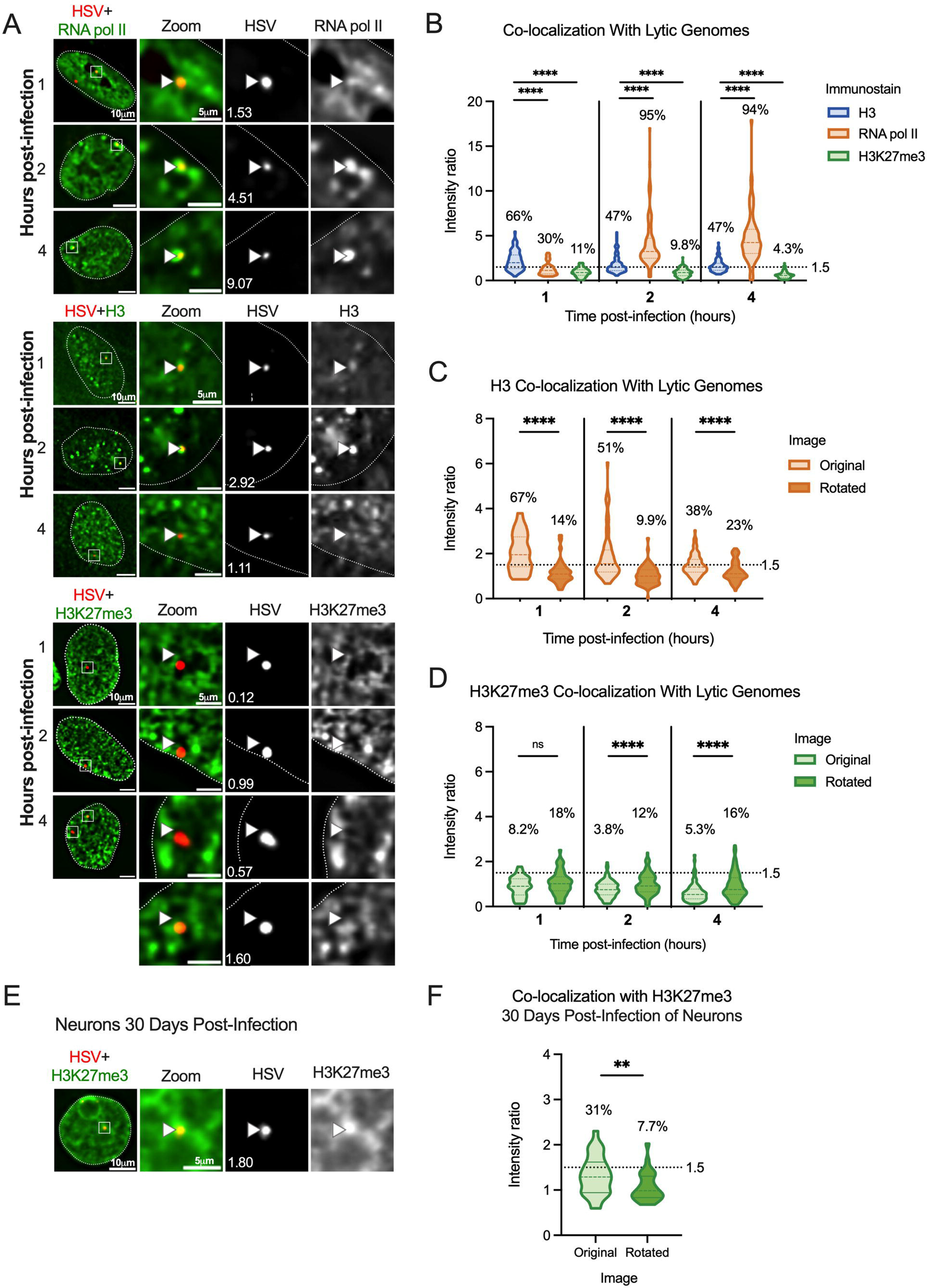
Incoming HSV-1 genomes do not co-localize with H3K27me3 during early lytic infection. HFFs were infected at an MOI of 3 PFU/cell with HSV^EdC^, fixed at different times post-infection and processed for click chemistry and immunostaining against H3K27me3. (A) Representative images of HFF nuclei 1, 2 and 4 hours post-infection, and zoomed image of individual viral genomes. NucSpotA intensity ratios are superimposed on each viral genome’s single channel image, with arrows for reference in the same spot in each channel. (B) NucSpotA quantification of image sets represented in A. Significance shown is based on the Kruskal-Wallis test. Each data point represents one viral genome. (C) Rotation analysis of co-localization with H3 at each time point as outlined in Figure 2. Paired analysis was performed for each genome (Wilcoxon test). (D) Rotation analysis for H3K27me3 intensity compared to those expected by chance (paired Wilcoxon test). Data shown in C and D were generated from the same original images quantified in B. (E) A representative image showing the nucleus of a latently infected neuron, within which a viral genome is co-localized with H3K27me3. (F) Rotation analysis of H3K27me3 co-localization with latent HSV genomes in neurons (paired Wilcoxon test). Percentages indicate the proportion of genomes with NucSpotA intensity ratios above the denoted co-localization threshold (dashed line). Adjusted p-values: **=<0.002, ****=<0.0001. N≥3.

We then used NucSpotA to quantify the enrichment of individual viral genome foci with H3K27me3 (Figure 3A, bottom). We observed a reduced association of viral genomes with H3K27me3 compared to total H3 at 1 hour post-infection (hpi), which was statistically significant (Figure 3B). In contrast, RNA polymerase II intensity ratios were even higher than for H3, resulting in the strongest co-localization with viral genomes at each time post-infection. Therefore, H3 and RNA polymerase strongly co-localized with viral DNA as expected, but H3K27me3 appears not to co-localize with lytic genomes based on the results of the overall NucSpotA analysis.

The above data indicate that, overall, viral genomes show reduced association with the H3K27me3 compared to host chromatin, and reduced levels also compared to total H3. However, this bulk NucSpotA analysis still did not take into account the possibility of a minority population of genomes that associate with H3K27me3.

Therefore, we set a cutoff (intensity ratio 1.5) above which genomes look visually co-localized with H3K27me3 when assessed qualitatively. The percentages of genomes above this cutoff (labeled as a dotted line on Figures 3B-D and 3F) serve as an indicator of whether a subpopulation of genomes co-localizes with H3K27me3. Notably, this method can be used to assess the heterogeneous association of viral genomes with any nuclear protein of interest. Using this method, we observed that 11%, 9.8%, and 4.3% of viral genome foci had enrichment values for H3K27me3 above this threshold at 1-, 2- and 4 hpi, respectively (Figure 3B). As a positive control for H3K27me3 co-localization, we also performed Click chemistry and immunostaining for latent genomes in mouse superior cervical ganglia (SCG) neurons (Figures 3E, 3F). In latently infected neurons, we observed enrichment of H3K27me3 on approximately 31% of latent HSV-1 genomes based on a NucSpotA intensity ratio above 1.5. This is consistent with previous observations that H3K27me3 is both enriched on the latent HSV genomes and its removal is important in reactivation (12, 15, 34, 35, 78–81). These data also highlight the potential heterogeneity in the epigenetic nature of latent HSV genomes, which may relate to different levels of expression of the latency-associated transcript between individual neurons or differences in sub-nuclear genome localization (82, 83). Importantly, these data support our use of intensity ratios to quantify co-localization between HSV-1 genomes and H3K27me3.

It was not clear from this analysis whether positively co-localizing genomes represent a true association, or if they are more co-localized than we would expect by chance (random placement of a genome in the nucleus). We thus performed an additional analysis, using each image to generate its control by rotating the viral genome channel 90 degrees relative to nuclear and histone stain channels (Figure 2B). This allows paired analysis between an original genome’s intensity ratio and that for its random placement within the same nucleus (84). Original image H3 co-localization was significantly greater than that for its random control image at each time point (Figure 3C), as is the co-localization of latent genomes with H3K27me3. However, co-localization with H3K27me3 in fibroblasts was similar to or below that expected by chance at all three time points post-infection (Figure 3D). In conclusion, assessing the co-localization of lytic viral genomes with histone modifications suggests that lytic genomes do not stably co-localize with H3K27me3, in contrast to co-localization with total histone H3.

### Inhibition of H3K27me3 deposition or removal does not alter its co-localization with HSV-1 genomes in fibroblasts

Although the rotation analysis suggested that a similar proportion of viral genomes could co-localize with H3K27me3 as those observed to co-localize by chance, it was still possible that a minority population is targeted for H3K27me3. Therefore, to thoroughly investigate whether this reflected deposition onto the viral genome, we pre-treated cells with UNC1999 (85), an inhibitor of the H3K27 methyltransferases EZH1 and EZH2 in the PRC2 complex, and quantified co-localization of viral genomes in the absence of H3K27 methyltransferase activity. Western blots of the total levels of H3K27me3 in fibroblasts demonstrated that 1.8 μM had the capability of reducing levels over time (Figures S2B). Therefore, we pre-treated cells with 1.8 μM UNC1999 followed by infection with HSV-1, maintaining drug treatment during and after infection. At 2 hpi, the co-localization of H3K27me3 at viral genomes was not significantly lower than for vehicle control (DMSO)-treated cells at the population level. The proportion of genomes above the intensity ratio threshold of 1.5 also did not decrease with PRC2 inhibition (Figure 4A). Therefore, we conclude that the small proportion of viral genomes with a threshold above 1.5 (5%) was not a result of active deposition of H3K27me3.

**Figure 4.**
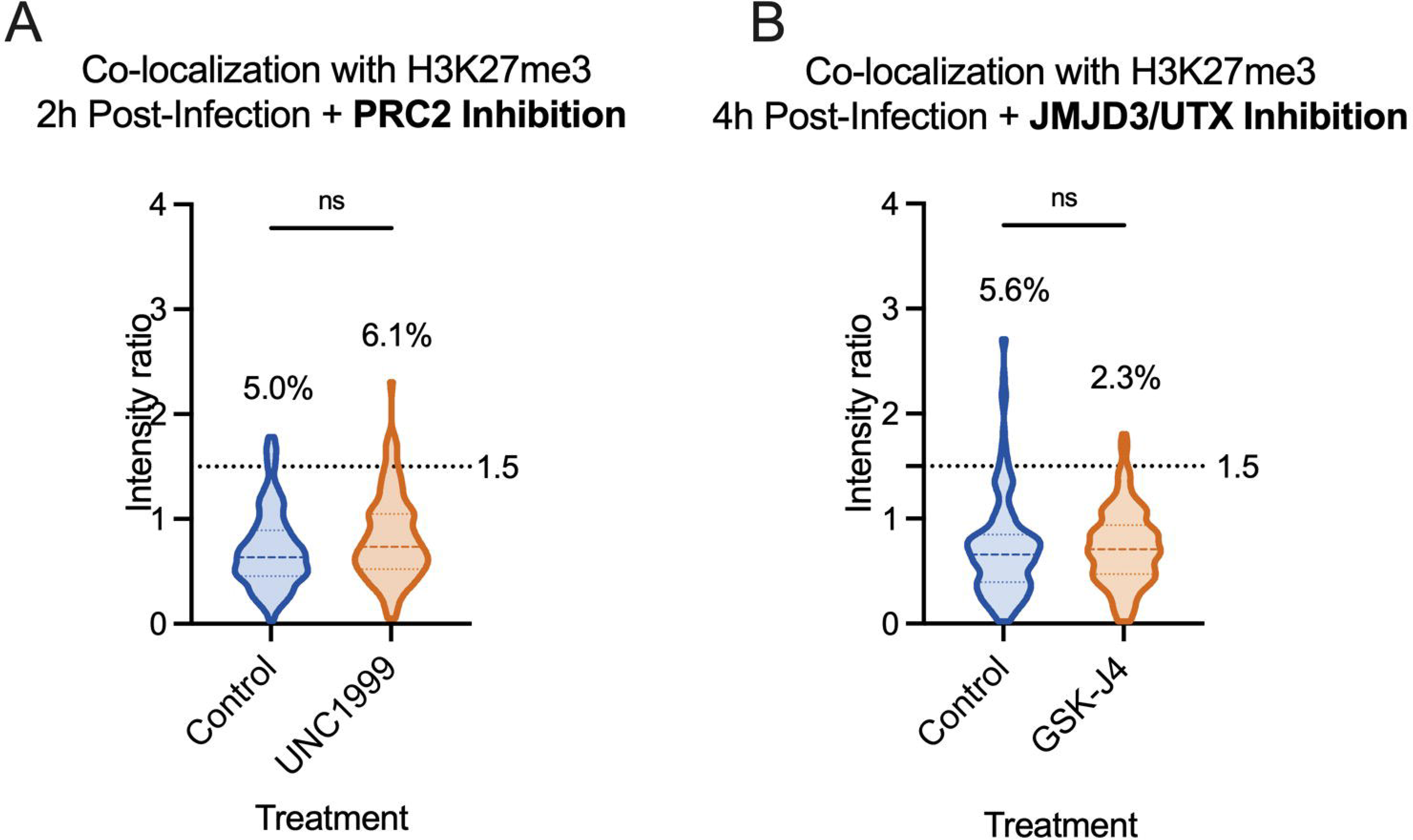
Inhibition of H3K27me3 dynamics does not impact H3K27me3 co-localization with lytic genomes. HFFs were infected with HSV^EdC^ following pre-treatment with inhibitor or vehicle control, maintaining treatment the throughout infection. (A) Colocalization of viral genomes with H3K27me3 at 2 hpi, treated with vehicle control or UNC1999. (B) Co-localization of viral genomes with H3K27me3 by 4 hpi, treated with vehicle control or GSK-J4 (10µM). Percentages represent genomes with co-localization above the threshold of 1.5. Kolmogorov-Smirnov tests, N≥3.

A final possibility was that H3K27me3 could be rapidly deposited and removed from viral genomes. Therefore, we added GSK-J4, an inhibitor of the H3K27me3 demethylases JMJD3 and UTX (86). Cells were again pre-treated with GSK-J4 (10 μM), and the inhibitor was included during the infection. Inhibitor activity at this concentration was confirmed by assessing H3K27me3 retention on cellular chromatin by western blot (Figure S2A). At 4 hpi, we did not observe an increase in the proportion of viral genomes co-localizing with H3K27me3 in the presence of the inhibitor (Figure 4B), suggesting the mark was not added and then rapidly removed by the activity of these histone demethylases. Taken together, these data suggest that H3K27me3 is not deposited on lytic genomes in HFFs during the early stages of lytic infection of fibroblasts.

### H3K27 demethylase inhibition restricts lytic gene expression

To determine whether the presence of methylated H3K27 can impact HSV gene expression in fibroblasts, we carried out gene expression analysis on cells infected in the presence of UNC1999. HFFs were again pre-treated with UNC1999, and then infected with HSV-1, with the inhibitor treatment maintained throughout infection. We then performed RT-qPCR to quantify lytic gene expression. We expected PRC2 inhibition with UNC1999 to enhance lytic gene expression if H3K27me3 were deposited on the viral genome. However, we did not observe any change in the expression of the IE mRNA *ICP27* and early mRNA *ICP8* transcripts with UNC1999 treatment (Figures 5A, 5B). A previous study found that long-term treatment with a high dose of UNC1999 can result in the expression of anti-viral genes including IL6, IFNA2 and IFNA1, and inhibition of viral gene expression (87). However, in our experiments using a lower dose of UNC1999 (3 μM) and a shorter time frame of treatment, we did not observe changes in *IL6* expression (Figure S2C), indicating the UNC1999 was not inducing an antiviral response in our cells that would otherwise impact the interpretation of these gene expression experiments. Therefore, these results indicate that the deposition of H3K27 methylation does not impact HSV-1 gene expression and support our conclusion that H3K27me3 is not being deposited on the HSV-1 genome during lytic infection.

**Figure 5.**
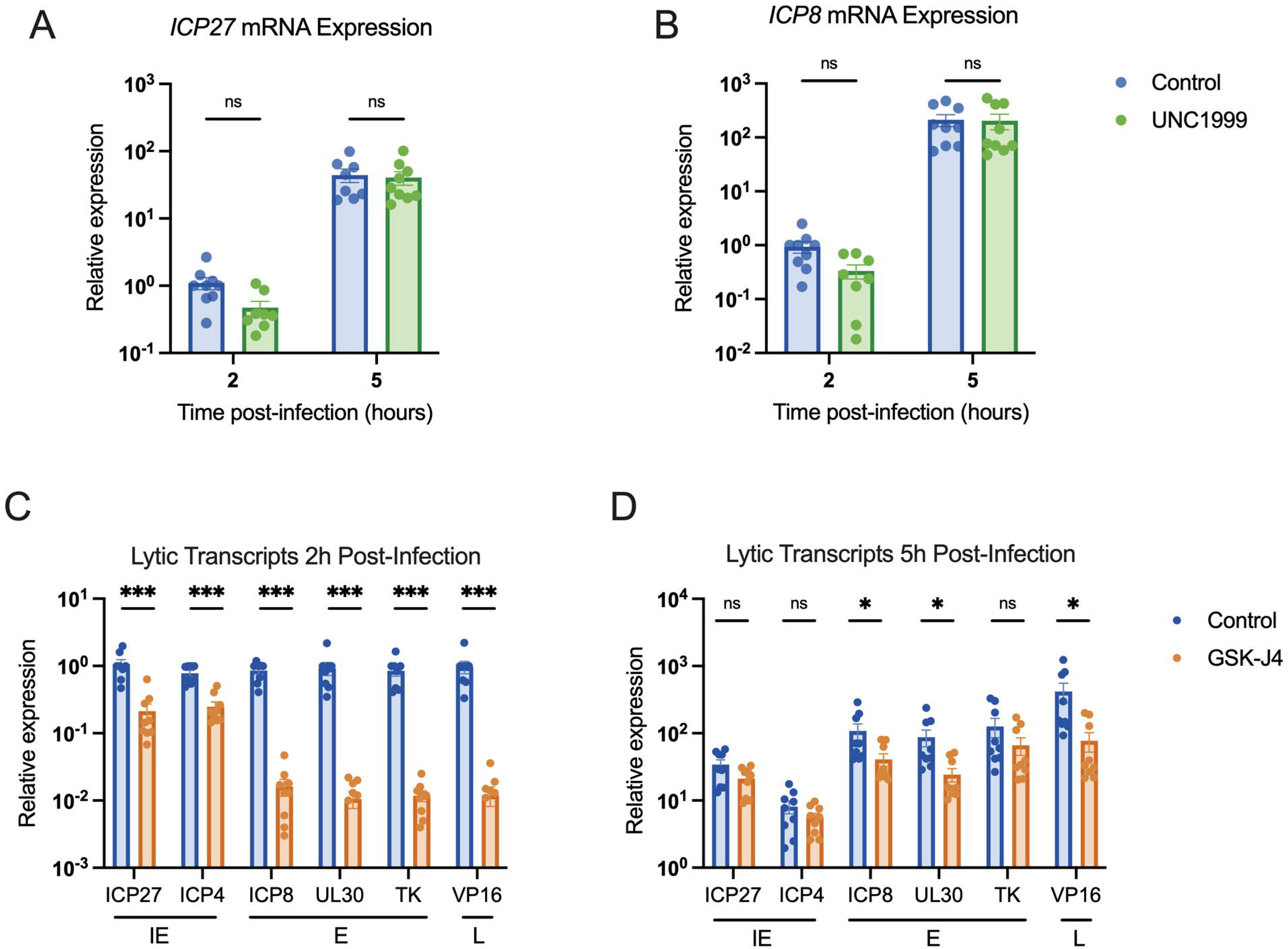
Inhibition of H3K27 demethylase activity restricts lytic gene expression, but inhibition of H3K27 methylation does not impact lytic gene transcription. HFFs pre-treated with inhibitor or vehicle control were infected at MOI 3 PFU/cell, maintaining treatment throughout infection. RNA lysate was harvested, cDNA synthesized and transcript levels determined by RT-qPCR. Fold expression change is relative to cellular g-actin transcript levels. (A) Relative levels of IE transcript ICP27, (B) and E transcript ICP8, in cells treated with vehicle control or UNC1999 (1.8µM). (C) Transcription of all three lytic gene classes (immediate early (IE), early (E) and late (L), as indicated) 2 hpi, vehicle control-treated or treated with GSK-J4 (10µM). (D) Transcription representative viral lytic genes 5 hpi. N N≥3; biological repetitions shown. Mann-Whitney test, adjusted p-values: *=<0.05, ***=<0.0005.

We also carried out the converse experiment and examined whether inhibition of the removal of H3K27 methylation impacted HSV-1 gene expression in fibroblasts using GSK-J4 (10 μM). Unexpectedly, we observed a repressive effect with GSK-J4 treatment. Although data up to this point suggest that H3K27me3 is not forming on the lytic genome, preventing H3K27 demethylation led to repression of all the immediate-early (IE), early and late transcripts checked at 2 hpi (approximately 3-5, 50-80 and 80-fold respectively; Figure 5C). By 5 hpi, some effect was still seen for early genes (2-3.5-fold), and leaky late gene *VP16* (5.4-fold), although this was less than that observed at 2 hpi, indicating the repression may be overcome later in infection (Figure 5D). Therefore, inhibition of JMJD3 and UTX activity limits, but does not fully prevent, HSV-1 lytic gene expression. This was surprising given our observation that inhibition of the H3K27 demethylases did not impact levels of H3K27me3 association. However, it was possible that inhibition of removal of other forms of H3K27 methylation would impact HSV-1 lytic gene expression.

### A subpopulation of genomes co-localize with H3K27me2 when H3K27 demethylation is inhibited

Given that the PRC2 and JMJD3/UTX complexes are responsible for methylation dynamics between all three methylation states of H3K27, we considered that another methylation state could be present and repressive to the lytic genome. We focused on H3K27me2, as this modification is also repressive to transcriptional activity (69, 70). Additionally, we considered that previous studies investigating the mechanisms of *de novo* Polycomb silencing in murine embryonic stem cells (mESCs) have shown full tri-methylation of H3K27 to take approximately 36 hours, a time frame inconsistent with the rapid events unfolding during early HSV-1 infection (67, 88, 89). However, the same studies demonstrate that H3K27me2 forms more rapidly. H3K27me2 is relatively understudied but is also associated with gene silencing. It protects against the deposition of H3K27 acetylation (an activating modification), and the H3K27me2 reader protein PHF20L1 has been shown to restrict transcription (69, 90). Notably, H3K27me2 is also one of the most abundant histone PTMs on the host genome and is more prevalent than H3K27me3 (69).

We therefore performed CUT&RUN and co-localization experiments, this time investigating the H3K27me2 modification. We found it difficult to source an H3K27me2 antibody with appropriate binding specificity and performed multiple experiments with one antibody (Diagenode C15410193) that turned out to have high binding affinity to unmodified histone H3 (Figure S1D). Histone peptide binding array analysis was conducted for four additional antibodies marketed to recognize H3K27me2, but none were selective for this mark (Figure S1B-E). A comparison of co-localization and CUT&RUN viral genome coverage using Diagenode C15410193 is shown in Figures S3A-B. We note although we cannot fully rule out that the CUT&RUN signal is from H3K27me2, the broad distribution across the genome pointed to non-specific binding. In addition, the co-localization of H3K27me2 with the viral genome using Diagenode C15410193 is higher than other H3K27me2 antibodies that were included in this study; therefore, we did not continue experiments using Diagenode C15410193. The explanation for the enhanced binding of this antibody to viral genomes is unclear but may result from non-specific binding to unmodified histones. We include the data here as an example of the need to accurately validate the binding specificities of histone antibodies.

We analyzed H3K27me2 using a more specific antibody as determined by histone peptide arrays (Figure S1B; Active motif 39245). Figure 6A shows representative images of an infected nucleus at each time point post-infection. Without any inhibitor treatment, the co-localization of viral DNA with H3K27me2 was below that for total H3 (Figure 6B), and similar to or below that expected by chance from rotation control analysis (Figure 6C). Further, PRC2 inhibition with UNC1999 (1.8 μM) did not reduce the co-localization of viral DNA with H3K27me2, indicating that under these conditions we either could not detect active deposition of H3K27me2 onto viral genomes or that it was rapidly removed (Figure 6E). Inhibition of H3K27 demethylation using GSK-J4 (10 μM) did cause a modest but significant increase in the fraction of viral genome foci that co-localize with H3K27me2; up to 15.5% of genomes (Figure 6F). Notably, the percentage increase in genomes co-localizing with H3K27me2 was reproducible between independent biological replicates, including biological repetitions with a separate H3K27me2 antibody (Active motif 61435 Figure S1E, Figure 6G). Therefore, these viral genomes (representative images Figure 6D) may represent a subpopulation of genomes that experience H3K27me2 deposition followed by removal by JMJD3 and/or UTX.

**Figure 6.**
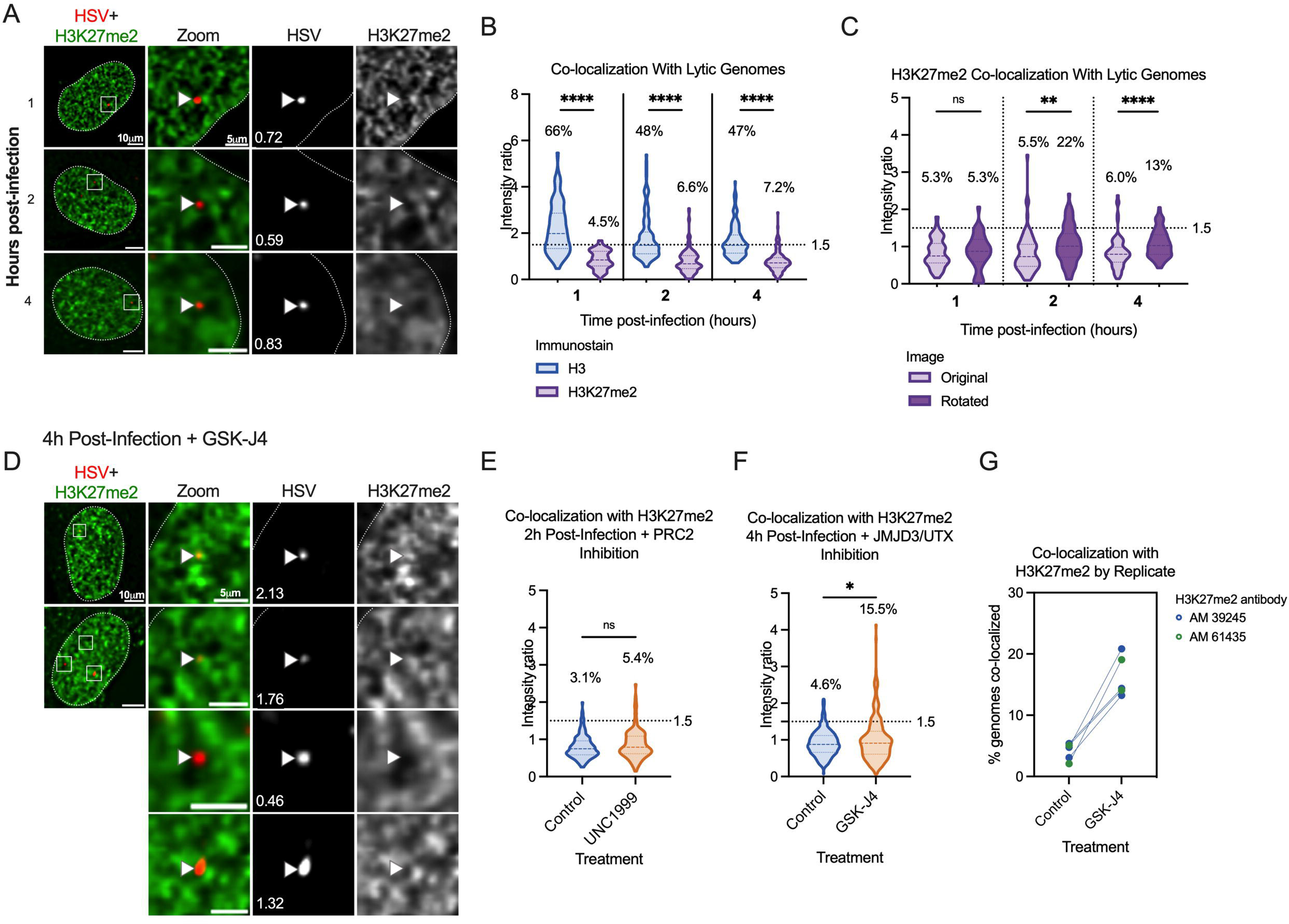
A sub-population of viral genomes co-localizes with H3K27me2 when H3K27 demethylation is inhibited. A) Representative images of HFF nuclei infected with HSV^EdC^ at 1, 2 and 4 hpi (Active Motif 39245 antibody). Intensity ratios are superimposed on each viral genome’s single channel image. (B) H3K27me2 compared to H3 co-localization with lytic genomes. (C) Rotation analysis for H3K27me2 images to compare actual co-localization to that expected by chance 1, 2 and 4 hpi. (D) Representative images of H3K27me2 co-localization with viral genomes in cells pre-treated and continuously treated with demethylase inhibitor GSK-J4 (10µM). (E) H3K27me2 co-localization with lytic genomes 2 hpi with vehicle control or UNC1999 treatment (1.8µM). (F) H3K27me2 co-localization with lytic genomes 4 hpi with vehicle control or GSK-J4 treatment (10µM). (G) Individual experimental replicates for GSK-J4 treated and vehicle control cells, including data included in F and two more data points with a different H3K27me2 antibody (Active motif 61435).

### CUT&RUN reveals a low association of lytic gene promoters with H3K27me2 that increases with K27 HDM inhibition

To investigate whether we could verify the association of H3K27me2 with the viral genome and any changes with GSK-J4 treatment, we performed CUT&RUN with paired-end sequencing, again using an antibody validated to bind H3K27me2 (CST D18C8 9728, Figure S1C). We were unable to find any previous studies investigating H3K27me2 association with the host genome in fibroblasts. Therefore, we found regions with high and low association in our dataset for comparison with the HSV-1 genome. We chose the *SERPINA1* promoter as a region of high enrichment and the *GAPDH* promoter as a region depleted for H3K27me2. In comparison to the *SERPINA1* promoter, we observed modest enrichment on viral promoters at 2 hpi. The enrichment of H3K27me2 was much lower at 4 hpi, likely because of ongoing viral DNA replication at this time-point, active removal of the modification, or of histone H3 itself, which has previously been reported independently of viral DNA replication (31). Notably, we did observe an increase in H3K27me2 levels on the host genome between 2- and 4-hours post-infection. Although a comparatively low level of H3K27me2 was detected on viral genomes at 2 hpi, Figure 7B shows the same enrichment values plotted without scaling to the host positive control. This representation shows an increase in H3K27me2 association in the presence of GSK-J4 at 2 hpi, indicating that a subpopulation of viral genomes may retain H3K27me2 in the presence of H3K27 demethylase inhibition.

**Figure 7.**
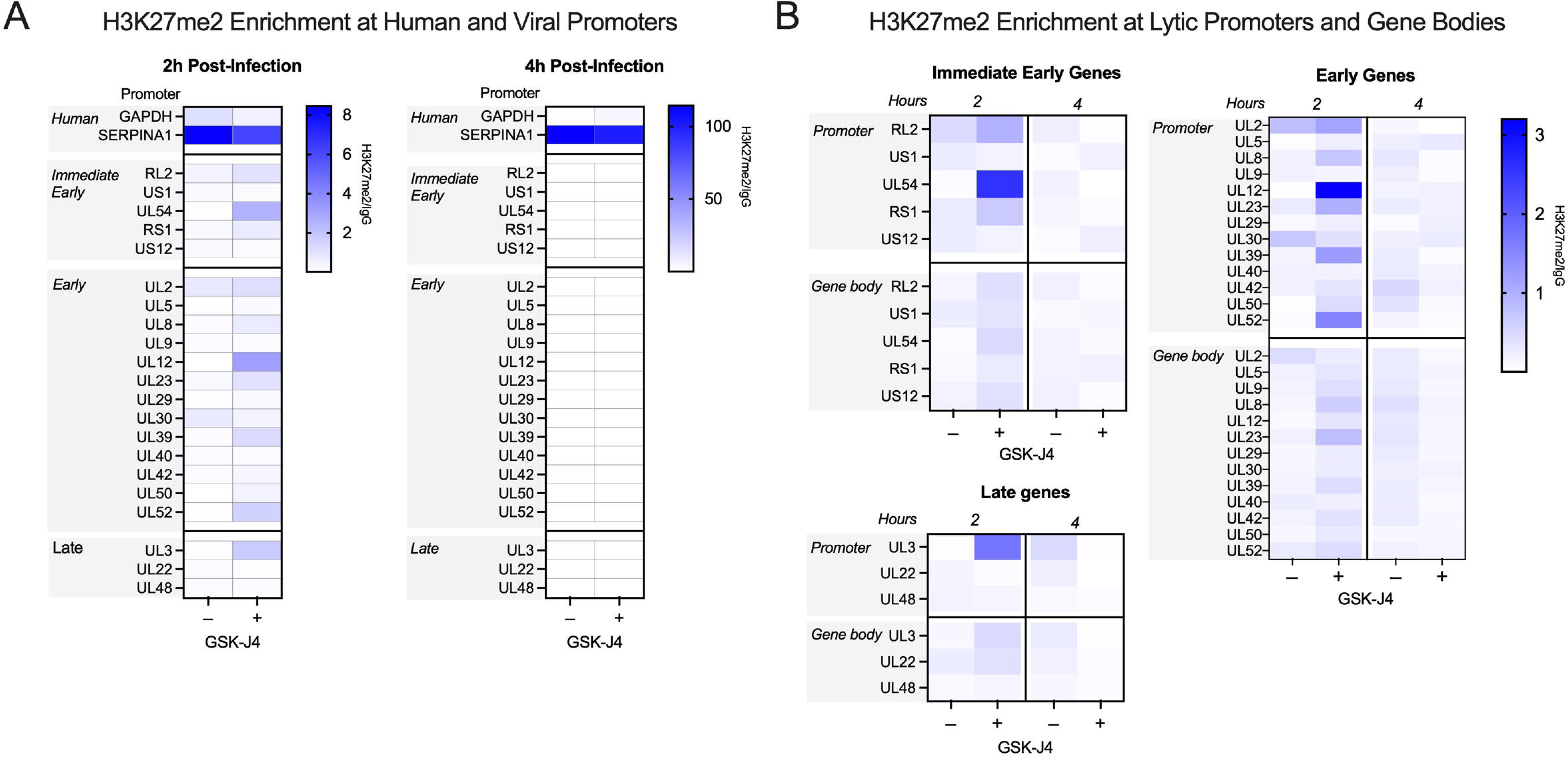
Bulk-level analysis of viral chromatin by CUT&RUN shows modest H3K27me2 enrichment at viral promoters, and less across gene bodies, during lytic infection. HFFs infected at MOI 3 PFU/cell, untreated or treated with 10µM GSK-J4, 2 and 4 hpi. The sum of coverage at defined promoter regions was used to calculate fold enrichment of H3K27me2 over IgG. (A) The geometric mean of two replicates’ fold enrichment is plotted as a heat map scaled to host gene *SERPINA1*. (B) The geometric mean of two replicates’ fold enrichment at viral promoters and gene bodies, scaled to viral enrichment only. CUT&RUN and downstream processing was carried out in parallel for two independent infections (N=2).

There appeared to be no correlation with gene class for the IE and early genes, although *UL54* (encoding ICP27) had the highest level of enrichment. Overall, the enrichment was higher in promoter regions compared to the gene bodies (Figure 7B). Taken together, these results indicate that H3K27me2 is deposited and removed on at least a subpopulation of viral genomes, and this removal enables more robust viral lytic gene expression at early times during infection.

### Transcriptionally repressed genomes are enriched for H3K27me2 in the absence of PML-NBs

We next explored whether we could enrich for viral genomes with the H3K27me2 modification, both to further validate that this modification is indeed targeted to viral genomes in fibroblasts and to determine under what conditions its deposition may occur. The methyltransferase activity of the PRC2 complex is known to be inhibited under conditions of active transcription (91, 92). Viral lytic gene expression is stimulated by the tegument protein, VP16, and the activation domain (AD) of VP16 recruits host proteins that promote transcription and limit total histone association (28). Therefore, taking into account this known function of the VP16AD, we investigated whether mutation of this domain resulted in increased H3K27me2 deposition.

We prepared EdC-labelled stocks of the previously described VP16AD mutant (RP5, KOS parent strain) (93). Initial parallel infections with RP5 compared to its rescued virus (RP5R) at an MOI of 3 PFU/cell showed a higher number of RP5 genomes reaching the nucleus than for RP5R, despite infection at the same PFU/cell; this is likely reflective of a reduced ability of RP5 to plaque on the U2OS cells used to grow and titer the viruses (data not shown). We thus adjusted the MOI of RP5 to achieve approximately 3 foci per nucleus when visualized with Click chemistry. The expected reduction in viral gene expression by RP5 compared to RP5R was confirmed by RT-qPCR from HFFs, although RP5 viral gene expression did still increase between 2 and 5 hpi albeit at a much-reduced level compared to the rescued virus (Figure 8A). We then analyzed the co-localization of RP5 genomes with H3K27me2 in HFF-telomerase immortalized cells (HFF-Ts) and observed approximately 24% that showed positive co-localization with H3K27me2. However, this was not significantly above the level for random placement determined using the rotated control images (Figure 8C).

**Figure 8.**
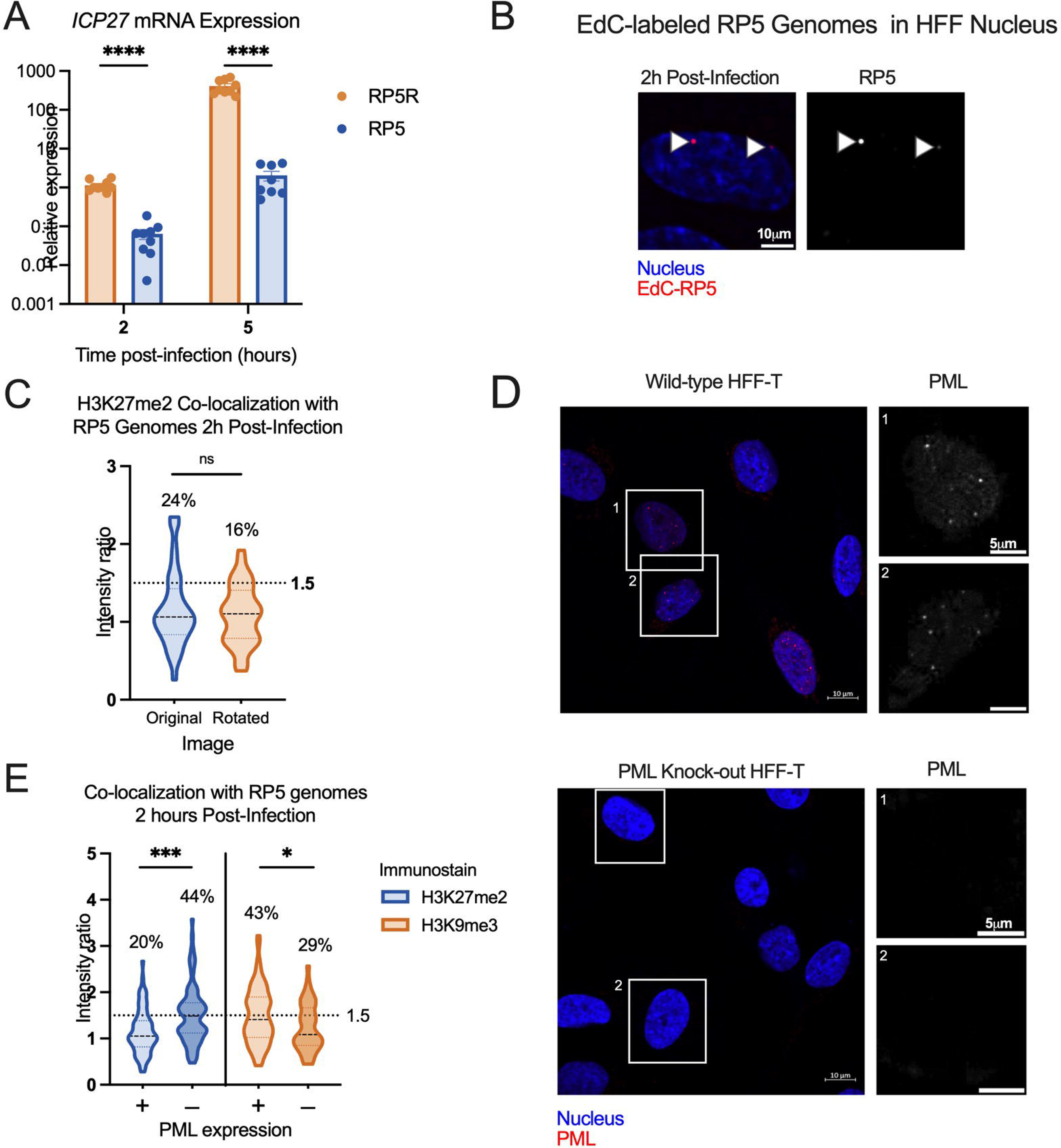
Transcriptionally repressed viral genome association with H3K27me2 is favored in the absence of PML expression. A) HFFs were infected at an MOI of 3 PFU/cell with either VP16 activation domain mutant RP5, or its rescue RP5R. Relative *ICP27* mRNA expression for RP5 infected cells, in comparison with RP5R-infected cells 2 and 5 hpi (multiple Mann-Whitney tests). Data are from 3 independent infections of 2-3 wells in parallel. (B) A representative image showing EdC-labeled RP5 genomes in an HFF nucleus 2 hpi. (C) H3K27me2 co-localization with RP5 genomes 2 hpi as determined using image rotation analysis (paired Wilcoxon test). (D) Confirmation of nanoblade-mediated PML knock-out, comparing wild-type and knock-out HFF-Ts immunostained for PML. Cells were clonally selected following nanoblade treatment. Panels in grayscale are zoomed in views of individual nuclei outlined in the left image. (E) H3K9me3 and H3K27me2 co-localization with RP5 genomes 2 hpi in the absence of PML, co-localization in PML-expressing cells compared with PML knock-out cells (Kolmogorov-Smirnov tests). Percentages indicate the proportion of genomes with NucSpotA intensity ratios above the denoted co-localization threshold (dashed line) in C and E. Data are from 3+ independent infections. Active Motif 39245 antibody was used for H3K27me2 immunostaining. Adjusted p-values: *=<0.05, ***=<0.0005., ****=<0.0001.

However, it has previously been reported in several studies that transcriptionally inactive HSV genomes associate with repressive PML-NBs in non-neuronal cells (61, 66, 82). PML-NBs can promote the deposition of H3K9me3 but are less linked to the deposition of H3K27me2/me3. Therefore, we asked if the presence of PML-NBs was preventing association with H3K27me2 on these transcriptionally repressed genomes by promoting the more constitutive H3K9me3 association. We created PML knock-out HFF-Ts (Figure 8D), the infection of which resulted in a significant increase in H3K27me2 association with RP5 at 2 hpi over PML-expressing HFF-Ts (wild-type HFF-Ts). Therefore, these data indicate that H3K27me2 associates with the HSV-1 genome either as a consequence of transcription repression and/or lack of the VP16AD in the absence of PML-NBs. Finally, to determine whether PML-NBs increase the association with constitutive heterochromatin, we also measured H3K9me3 co-localization with RP6 genomes. We found an increase in H3K9me3 association at 2 hpi in the wild-type HFF-Ts (Figure 8E), indicating that PML knock-out cells indeed favor H3K27me2 formation over H3K9me3 formation at transcriptionally inactive viral genomes.

### Association of the H3K27me2 reader protein PHF20L1 with a sub-population of HSV-1 genomes

To explore the repressive functional outcome of H3K27me2 formation on lytic HSV-1 genomes, we investigated the co-localization of viral genomes with a reader of this histone PTM. PHD Finger Protein 20-Like Protein 1 (PHF20L1) was reported as an H3K27me2 reader in the context of breast tumor growth, working with PRC2 and nucleosome remodeling and deacetylase (NuRD) complexes to facilitate transcriptional repression (70). PHF20L1 co-localization was consistently found in approximately one third of genomes at 4 hpi (34% of genomes), and an increase in co-localized genomes was observed with GSK-J4 treatment. This increase is reflected in a 9% increase in genomes above the co-localization cutoff, as well as a statistically significant difference between the two data sets (Figure 9B). Notably, the 9% increase in genomes co-localizing with PHF20L1 is similar to the 11% increase in co-localization with H3K27me2 (Figure 6F-G.) Representative images in the presence and absence of GSK-J4 are shown, including one genome that is co-localized with PHF20L1 (intensity ratio 1.72) and one that is not (intensity ratio 0.69) within the same nucleus (Figure 9A).

**Figure 9.**
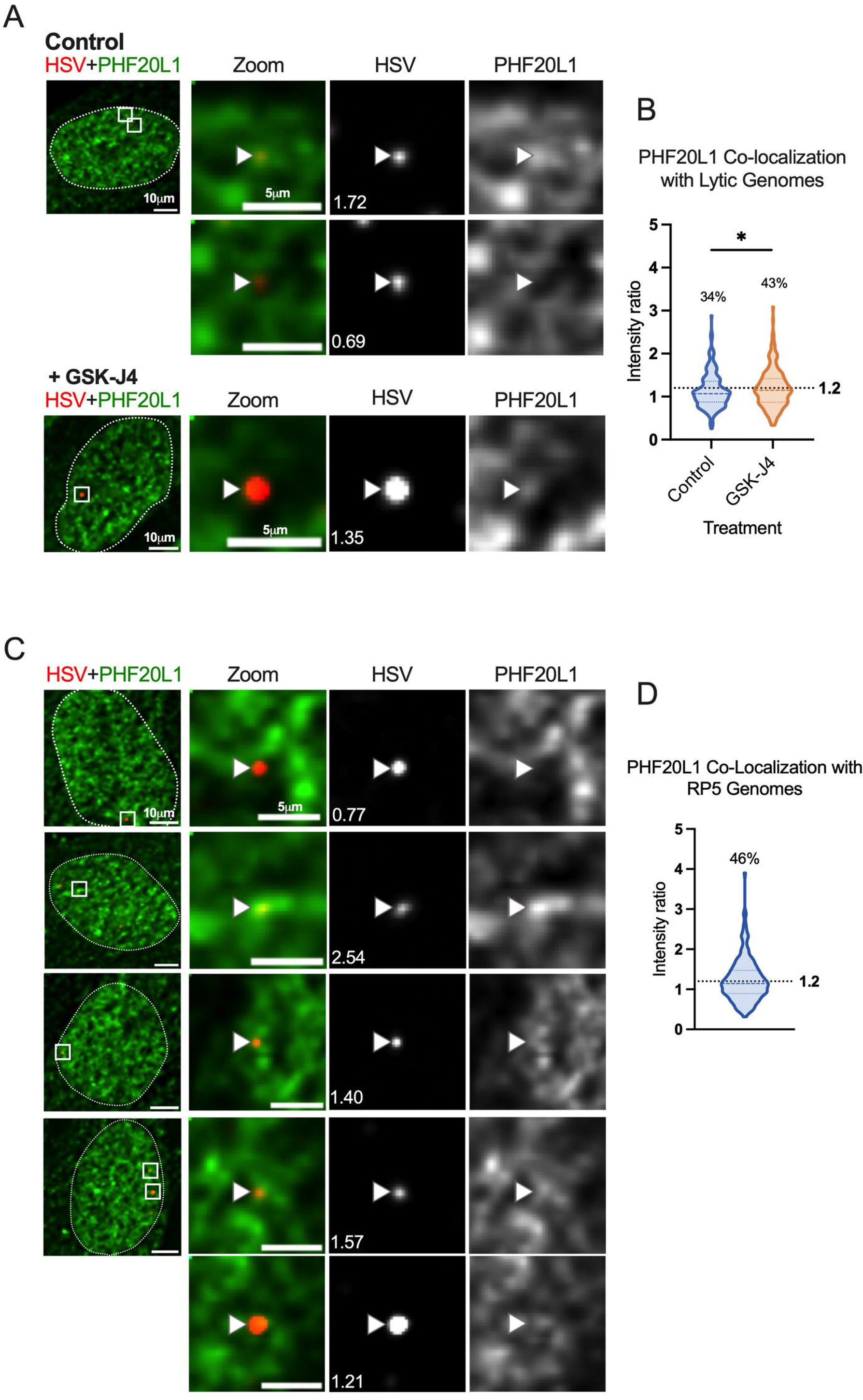
H3K27me2 reader PHF20L1 co-localizes with a sub-population of lytic genomes, including transcriptionally repressed genomes in the absence of PML expression. A) Representative images of HFFs infected with HSV^EdC^ immunostained for PHF20L1 in both control and GSK-J4 (10µM) treated conditions 4 hpi. Cells were pre-treated for 2 hours before infection at an MOI of 3 PFU/cell, and treatment maintained throughout infection. (B) Quantification of images represented in A, showing NucSpotA intensity ratios for PHF20L1 co-localization (Kolmogorov-Smirnov test) in vehicle control or GSK-J4-treated (10µM) cells. (C) Representative images showing PHF20L1 co-localization with transcriptionally inactive RP5 genomes in the absence of PML expression 4 hpi. PML knock-out HFF-Ts were infected with EdC-labeled RP5 to approximately 3 genomes per nucleus. (D) Quantification of RP5 co-localization with PHF20L1 in PML knock-out HFF-Ts 4 hpi. Intensity ratios are superimposed on each viral genome’s single channel image in A and C. Percentages indicate the proportion of genomes with NucSpotA intensity ratios above the denoted co-localization threshold (dashed line).

Because we observed the highest levels of H3K27me2 association with RP5 genomes in PML knock-out cells, we also investigated the association of RP5 genomes with PHF20L1 in these same cells. Co-localization was indeed seen in a large proportion of RP5 genomes (46%) 4 hpi, as shown with representative images (Figure 9C-D). This further validates the potential for transcriptionally inactive genomes without PML-NBs to be targeted for H3K27me2 and read by PHF20L1.

## Discussion

The process of lytic infection with HSV-1 is often described as a battle between the host cell and the infecting virus, with the host cell trying to silence gene expression by depositing repressive heterochromatin on the viral genome and the virus overcoming this silencing for gene expression to occur. However, for the Polycomb-associated modification, H3K27me3, there was little experimental evidence to support this model. Hence, using multiple techniques, we set out to determine whether any H3K27me3 could be detectibly deposited on the incoming genome by the host, which the virus removes for lytic replication to take place. This was important to understand, first, to determine how the host cell attempts to silence incoming foreign HSV-1 DNA, and second, because this modification is ultimately enriched on the HSV-1 genome during a latent infection of neurons (12, 15, 19, 20, 34, 35). Although we were unable to detect H3K27me3 enrichment on lytic genomes, are data suggest that a subpopulation of genomes are targeted for silencing by H3K27me2. This is intriguing because less is known about H3K27me2 versus the more commonly studied H3K27me3. The recent identification of a protein that specifically reads H3K27me2, PHF20L1, has illuminated a direct role for this modification in the recruitment of transcriptional repressors (70). In line with a role for H3K27me2 in lytic gene repression, we could also detect co-localization of viral genomes with PHF20L1 during lytic infection. Therefore, the deposition of H3K27me2 appears to play a more prominent role than H3K27me3 in repressing HSV-1 gene expression during lytic replication.

There are some caveats to our study. One being that we used small molecule inhibitors to this histone methyltransferases and histone demethylases and have not carried out knock-out or knock-down experiments. However, in attempting to do these experiments, we found that knockdown of the histone demethylases resulted in enhanced lytic gene expression, which is consistent with previous studies showing that a that lack of these enzymes results in a reduction in anti-viral gene expression (86). Therefore, long-term loss of these enzymes in fibroblasts likely results in more indirect impacts on gene expression, making this approach a challenge. A further caveat is that antibody binding to histone PTMs can be influenced by neighboring PTMs. For example, binding to H3K27me2/me3 can be occluded by phosphorylation of H3S28. In preliminary studies using an H3K27me3/pS28 antibody, we were unable to detect co-localization with the viral genome. However, there are currently no antibodies that recognize the dual H3K27me2/pS28 modification state and therefore we cannot rule out the possibility that a subpopulation has these combined modifications.

Determining the presence or absence of a particular protein or histone PTM on viral genomes can be a challenge. Here, we decided to characterize the H3K27me2/me3 distribution across the early lytic HSV-1 genome by CUT&RUN using histone antibodies analyzed for their binding specificity (94). Using human genome loci to validate each antibody’s DNA yield, we observed a relatively low level of H3K27me2/me3 across the viral genome. However, because of the known heterogeneity in the fate of HSV genomes following infection of fibroblasts (43, 44, 46, 95–98), we developed a novel method for quantification of individual HSV genome foci with nuclear proteins. This is particularly important since comparison to host regions following CUT&RUN may obscure the enrichment at sub-populations of HSV genomes. NucSpotA analysis accounts for variability amongst a set of images by thresholding the green signal above a chosen percentage of its maximum intensity. We visually determined an intensity ratio above which co-localization is occurring but found that it is important not to rely on the number’s magnitude alone. Appropriate controls, such as a known positive control for co-localization or random placement in the nucleus through channel rotation, helped inform our interpretation of the data. Additionally, statistical analysis of the dataset applies to the whole population of genomes, but the shape of the violin plot (data point density) can be indicative of a sub-population of genomes emerging. This assay was particularly informative in combination with other variables, such as inhibition of H3K27 methylation dynamics and the use of PML knockout cells. In the future, we expect NucSpotA to be a useful tool for the field to analyze viral genome co-localization with nuclear proteins where heterogeneity in the fate of viral genomes can occur.

Our data suggests that H3K27me2 is deposited and removed from a subpopulation of lytic genomes. The combined data using H3K27 demethylase inhibitors and analyzing H3K27me2 genome co-localization and viral mRNA levels supports a model by which H3K27me2 is repressive to lytic gene expression from a subset of viral genomes and is removed to permit more robust expression. The full spectrum of factors that regulate the deposition of H3K27me2 onto a subset of genomes is not clear. Our data point to differences in the subnuclear positioning of viral genomes and potentially viral transcriptional activity. Additional factors, which need not be mutually exclusive include the cell state, stage in mitosis, diversity in viral genome sequence, amount of infecting virus and responses from neighboring abortively or fully infected cells, which have all been linked to heterogenous outcomes of HSV infection (43, 44, 46, 95–98).

The H3K27me2 modification is understudied compared to H3K27me3 despite it being a predominant modification on the host genome and also being implicated in the repression of gene expression (19). There are several important implications of this observation. The first would be the mechanisms of targeting of H3K27me2 to the viral genome and investigating whether this is consistent between neurons and non-neuronal cells. The PRC2 complex can be targeted and activated to methylate H3K27 via different mechanisms (19). One model for H3K27me2/me3 involves general targeting to chromatin, potentially via RNA or binding to unmethylated CpG motifs, and inhibition by single-stranded RNA and activating histone PTMs (99–101). Our data showing increased deposition onto the VP16 mutant virus potentially supports these mechanisms, however, we cannot rule out other direct roles of VP16 in inhibiting PRC2 recruitment or inhibition. PRC2 can also be recruited following ubiquitination of lysine 119 on histone H2A (H2AK119ub) by PRC1 (102, 103). However, this pathway has only been described in pluripotent cells and the protein that links H2AK119ub to PRC2 recruitment may not even be present in more differentiated cells (104). Further, it is unknown whether the HSV-1 genome is enriched in H2AK119ub in non-neuronal cells. However, as part of ongoing studies in our lab, we have found enrichment of H2AK119ub on latent viral genomes (Dochnal et al, unpublished).

Previous studies have found that PML-NBs can promote H3K9me3, but not H3K27me3 (32). Therefore, our data showing that PML-NBs may also limit H3K27me2 is consistent with this model. Notably, we previously found that primary neurons are devoid of PML-NBs but can form with type I interferon treatment (66). In addition, targeting of viral genomes to PML-NBs only occurred with type I interferon exposure. Therefore, by combining data from our previous study with our new data in fibroblasts, we can start to assemble a model by which the heterogeneity in the epigenetic structure of the latent genome arises. For those genomes that enter neurons exposed to type I interferon, and likely exposure specifically on the soma (64, 66, 105, 106), would result in association with PML-NBs and H3K9me3 enrichment. For genomes that were not targeted to PML-NB, they would be more likely to become enriched for H3K27me2/me3. However, this model is based on extrapolating our findings here to neurons and therefore requires additional testing using relevant neuronal model systems.

Our findings that H3K27me2 can be deposited on incoming HSV-1 genomes but not H3K27me3 are consistent with our understanding of *de novo* H3K27me3 dynamics; reintroducing PRC2 activity leads to nucleation of H3K27me3 sites after 12 hours, and propagation across a region at 36 hours (67, 88, 89). However, these previous observations were made in undifferentiated, mouse embryonic cells, and we therefore had to consider that H3K27 methylation dynamics could be faster in the context of infecting a differentiated cell type. The factors that regulate the progression from H3K27me2 to H3K27me3 are not known. In the context of HSV latency establishment, this will be important to understand. A previous study showed that H3K27me3 did not form on latent genomes until 10-14 days post-infection of mice (12). Whether H3K27me2 forms prior to this and plays a role in lytic-gene repression during entry into latency is unknown.

Given the identified role for PHF20L1 as a repressive H3K27me2 reader, its co-localization with HSV-1 genomes strengthens the evidence for a mechanism by which H3K27me2 represses lytic genes soon after infection of a fibroblast. Hou et al. propose a model whereby PRC2 and NuRD complexes are recruited by PHF20L1 binding to H3K27me2, resulting in transcriptional repression in the context of breast tumorigenesis (70). This model may link H3K27me2 on lytic HSV genomes with the transcriptional repression we observed with H3K27 demethylase inhibition. Interestingly, proteomic studies have shown NuRD complex components associated with both input and replicating HSV-1 genomes (77, 107). This axis would represent a previously undescribed defense mechanism against foreign DNA, and a corresponding pro-viral role for H3K27 demethylation during lytic infection. It also remains to be determined whether PHF20L1 plays a role during the establishment of latency in a neuron, where it could be an important component of the factors regulating cell type-specific transcriptional outcomes of HSV-1 infection.

## Materials and Methods

### Cells, viruses and drugs

Primary human foreskin fibroblast (HFF), U2OS, and Vero cells were all obtained from American Type Culture Collection. Telomerase immortalized retinol pigmented epithelial cells (RPE-T) have been described previously (59). HFF-Ts were generated by telomerase immortalization by lentiviral transduction of HFFs using pLV-hTERT-IRES-hygro (a gift from Tobias Meyer (Addgene plasmid # 85140 ; http://n2t.net/addgene:85140 ; RRID:Addgene_85140)) (108). HFF, U2OS, HFF-T and RPE-T cells were cultured in Dulbecco modified Eagle Medium High Glucose (DMEM, Gibco 11965-092) supplemented with 10% fetal bovine serum (FBS). RPE-Ts were cultured in the presence of 5 μg/ml of Hygromycin. Vero cells were cultured in DMEM supplemented with 10% Fetalplex (GeminiBio 50-753-2987). 293-LTV cells (Cell Biolabs LTV-100) were cultured in DMEM High Glucose with 10% FBS and 1% MEM NEAA (Gibco 11140-050).

Herpes simplex virus (HSV-1) strain 17Syn+ was grown on Vero cells infected at an MOI of 0.1 PFU/cell, and cultured at 34°C for 2 days or until cytopathic effect was observed. Heparin sodium (BP2425, Fisher Scientific) in phosphate buffered saline (PBS) was added to flasks to a final concentration of 50µg/ml and incubated at 37°C for 4-6 hours, before supernatant collection and centrifugation at 4°C 150 RCF for 10 minutes. Supernatant was centrifuged at 20,000 RCF for 1 hour at 4°C, and virus pellet resuspended in 10% glycerol in PBS, then sonicated at amplitude 20% for 20 seconds before aliquoting and storage at −80°C. Stocks of 17Syn+ were titrated on Vero cells. VP16 activation domain mutant (RP5) and rescue (RP5R) HSV-1 stocks were generated by infecting U2OS cells at an MOI of 0.05 PFU/cell. RP5R and RP5 were titrated on U2OS cells as described previously (75). RP5 was propagated and titrated in the presence of hexamethylene bis-acetamide (HMBA) in PBS (Sigma Aldrich 224235-50G) to a final concentration of 3 mM.

Inhibitors were resuspended in dimethyl sulfoxide (DMSO) and vehicle controls performed with inhibitor-equivalent volumes of DMSO. UNC1999 (Cayman Chemical Company 14621) was used at 1.8µM, and GSK-J4 (Sigma Aldrich SML0701) used at 10µM unless otherwise indicated.

### PML nanoblade and knock-out cell production

Nanoblades were generated as previously described (109) in 293-LTV cells transfected using jetPRIME (Polypus 101000027) with plasmids pCMV-VSV-G (a gift from Bob Weinberg Addgene plasmid # 8454 ; http://n2t.net/addgene:8454 ; RRID:Addgene_8454), BaEVRLess (gifted by Els Verhoeyen; constructed with pCMV-VSV-G)(110), p5349 (pBS-CMV-gagpol, a gift from Patrick Salmon (Addgene plasmid # 35614 ; http://n2t.net/addgene:35614 ; RRID:Addgene_35614)), BIC-Gag-Cas9 (a gift from Philippe Mangeot & Théophile Ohlmann & Emiliano Ricci (Addgene plasmid # 119942 ; http://n2t.net/addgene:119942 ; RRID:Addgene_119942)(109) and pBLADE PML (target sequence GCG GGT GTG TCT GCA CCT AGG GG) or pBLADE non-targeted control (target sequence ATC GTT TCC GCT TAA CGG CG) (pBLADE constructs a gift from Chris Boutell, BLADE was a gift from Philippe Mangeot & Théophile Ohlmann & Emiliano Ricci (Addgene plasmid # 134912 ; http://n2t.net/addgene:134912 ; RRID:Addgene_134912) (109). Cas9 production was quantified by serial dilution as described using Cas9 nuclease (New Engliand BioLabs M0386S). HFF-hTERT cells were transduced with Nanoblades with 8 μg/mL polybrene (Boston BioProducts BM-862M-1) and cultured with 5 ng/ml human fibroblast growth factor (FGF, Gemini Bio-Products 300-113P) throughout clonal selection. Knock-out was verified by immunostaining for PML.

### Production of EdC-labeled virus stocks

The EdC-labeling protocol was adapted from previously published procedure (59, 60). RPE-Ts were infected with 17Syn+ in 0.2% FBS DMEM at an MOI of 0.001 PFU/cell for WT 17Syn+ and RP5R, or 0.5 PFU/cell for RP5 and kept at 33°C. HMBA was included in media at a final concentration of 3mM during EdC-labeling of RP5. EdC (Sigma-Aldrich T511307) pulses diluted in 0.2% FBS DMEM were added at 6-24h, 48h, and 72h post-infection to 1µM final concentrations. At 72 hpi heparin sodium was added, incubated and supernatant was passed through a 0.45 µm PES syringe filter and the supernatant virus harvested as above with the inclusion of an additional two wash steps using DMEM containing 0.2% FBS.

### Virus infections

Cells were plated into 24 well plates 24 h prior to infection. Where specified, cells were pre-treated for 1-2 h prior to infection with inhibitors. Virus was diluted in phosphate-buffered saline (PBS) containing 0.1% glucose and 1% FBS. In all experiments, this represents the 0 h time-point post-infection. After 1 h of adsorption at 37°C, cells were washed twice with PBS containing 0.1% glucose and 1% FBS. Cells were overlaid with DMEM containing 1% FBS and incubated at 37°C.

### Primary neuronal cultures

Sympathetic neurons from the superior cervical ganglia (SCG) of post-natal day 0–2 (P0-P2) CD1 Mice (Charles River Laboratories) were dissected as previously described (79). Rodent handling and husbandry were carried out under animal protocols approved by the Animal Care and Use Committee of the University of Virginia (UVA). Ganglia were briefly kept in Leibovitz’s L-15 media with 2.05 mM L-Glutamine before dissociation in Collagenase Type IV (1 mg/mL) followed by Trypsin (2.5 mg/mL) for 20 min each at 37°C. Dissociated ganglia were triturated, and approximately 5,000 neurons per well were plated onto rat tail collagen coated glass coverslips. Sympathetic neurons were maintained in CM1 (Neurobasal Medium supplemented with PRIME-XV IS21 Neuronal Supplement (Irvine Scientific), 50 ng/mL Mouse NGF 2.5S, 2 mM L-Glutamine, and Primocin). Aphidicolin (3.3 mg/mL) was added to the CM1 for the first 5 days post-dissection.

### Establishment of latent HSV-1 infection in primary neurons

Neonatal SCGs were infected at postnatal days 6-8 with EdC labelled HSV at an MOI of 7.5 PFU/cell assuming 5,000 cells/well in PBS supplemented with 1% FBS, 4.5 g/L glucose and 10 mM Acyclovir (ACV) for 3 hr at 37°C. Post-infection, the inoculum was replaced with CM1 containing 50 mM ACV.

### Click chemistry and immunofluorescence

Cells were plated onto glass coverslips in 24 well plates and infected at the indicated MOIs. Cells were washed twice with cytoskeletal (CSK) buffer (10 mM HEPES, 100 mM NaCl, 300 mM Sucrose, 3 mM MgCl2, 5 mM EGTA) then simultaneously fixed and permeabilized in 1.8% formaldehyde (methanol-free, Thermo Fisher Scientific 28906) and 0.5% Triton-X100 in CSK for 10 min. Cells were washed three times in PBS and twice in CSK post-fixation. Coverslips were blocked with 3% Bovine Serum Albumin (BSA, Fisher Bioreagents BP1600-100) prior to Click-chemistry followed by immunostaining. EdC-labelled HSV was detected using the Click-iT Plus EdU Alexa Fluor 555 Imaging Kit (Thermo Fisher Scientific C10638) according to the manufacturer’s instructions, using a working stock of picoyl azide-Alexa Fluor 555 (PCA-AF 555). For immunostaining, samples were incubated overnight with primary antibodies in 3% BSA and washed in PBS three times. Following primary antibody treatment, coverslips were incubated for one hour in Alexa Fluor conjugated secondary antibodies (Invitrogen A-11008). Nuclei were stained with Hoechst 33258 (Life Technologies H3570). Antibodies and their applications are listed in Table S1.

### Image analysis

Epifluorescence microscopy images were acquired at 60× using an sCMOS charge-coupled device camera (pco.edge) mounted on a Nikon Eclipse Ti Inverted Epifluorescent microscope using NIS-Elements software (Nikon). Individual nuclei were isolated from this field of view and 3D deconvolved using the Landweber method (10 iterations) in NIS-Elements software.

Deconvolved z-stacks processed by NucSpotA, which is part of the Mitogenie suite, using thresholds to isolate positive immunostaining signal (visually determined) as follows: 70.5% (all histone stains) and 65% (PHF20L1) for 17Syn+ infection of HFFs; 75% (H3K27me2), 70% (H3K9me3 and PHF20L1) for RP5 infection of HFF-Ts and PML knockout HFF-Ts. Rotation control images were generated from original channel combination images using FIJI, prior to analysis with NucSpotA. The stated co-localization thresholds were blindly calibrated by eye.

### Quantification of viral gene expression

Analysis of mRNA expression by reverse-transcription quantitative PCR (RT-qPCR). To analyze HSV mRNA relative expression, total RNA was extracted using the Zymo Research Quick-RNA MiniPrep Kit (R1055) with an on-column DNAse digestion. Reverse transcription was carried out on equivalent amounts of RNA using Maxima Maxima First Strand cDNA Synthesis Kit (Thermo Scientific K1642), RiboLock RNAse Inhibitor (Thermo Scientific EO0382), Random Hexamer Primer (Thermo Scientific SO142) and dNTP Set (Thermo Scientific R0181), and qPCR was carried out using PowerUp SYBR Green Master Mix (Applied Biosystems A25741). The relative mRNA copy number was determined using the 2^-ΔΔCt^ method and viral mRNAs were normalized to that of the human reference gene mRNA transcript from *ACTG1* (actin gamma 1). All samples were run in duplicate on an Applied Biosystems Quantstudio 6 Flex Real-Time PCR System and analysis carried out using QuantStudio Real-Time PCR Software v1.7. Primer sequences: ICP8 as published (78); TK as published (111); others are listed in Table S2.

### Western blotting

Confluent HFFs cultured in a 6-well plate were treated with indicated concentrations of GSK-J4 or UNC1999 in 10% FBS DMEM for four days, with a media change including fresh inhibitor on day two. Untreated cells were cultured in parallel. Histones were isolated from inhibitor treated or untreated cells using the histone extraction kit (Active Motif, 40028) and western blots performed. Histone extracts were combined with Li-cor 4X Protein Loading Buffer (928–40004) and resolved on Bio-Rad Mini-PROTEAN TGX 4-20% gel (4561094) in Boston BioProducts Tris-Glycine-SDS Running Buffer (BP-150), and transferred onto an Immobilon-FL PVDF membrane (IPFL00010) using Boston BioProducts Transfer Buffer (BP-190) made to 20% v/v methanol. Membranes were blocked in Odyssey Blocking Buffer (OBB, 10 mM Tris-HCl pH 7.5, 150 mM NaCl, 2% Fish gelatin, 1% Ovoalbumin) for a minimum of 1 hour at room temperature, washed with TBS-T (Research Products International T60075-4000.0 with 0.1% Tween 20, and incubated with primary antibody (diluted in OBB with 0.2% Tween 20) overnight at 4°C. Li-cor secondary antibodies (925–32211, 925–68070) were diluted in OBB (with 0.2% Tween 20, 0.02% SDS) and blots imaged on a Li-Cor Odyssey CLX-1374 . Band intensity quantification was performed using Li-Cor Image Studio v5.2. Li-Cor Chameleon Duo Pre-Stained Protein Ladder (928–60000) was used.

### Histone peptide array

Peptide synthesis and validation, array fabrication and antibody analysis were performed as described (42, 112–114). Each peptide was spotted in triplicate twice per array. Triplicate spots were averaged and treated as a single value for subsequent statistical analysis as described (115).

### CUT&RUN

CUT&RUN was carried out using the Epicypher CUTANA Chic/CUT&RUN Kit and workflow (14–1048). Antibodies used for CUT&RUN are included in Table S1. Dual indexed DNA libraries were prepared using Epicypher CUTANA CUT&RUN Library Prep Kit (14–1001). Pair-ended, partial lane sequencing and de-multiplexing was carried out using NovaSeq (Novogene). Data analysis was performed using command line and R code, and workflow, adapted from the cited tutorial (116). The Rivanna high-performance computing environment (UVA Research Computing) was used for command line data processing.

The HSV-1 Syn17+ genome sequence (NCBI NC_001806.2) was used in combination with the hg38 human genome assembly (RefSeq GCF_000001405.40) to make a joint Bowtie2 index genome. Sequence alignment was performed with bowtie2 with the following settings*: --end-to-end --very-sensitive --no-mixed --no-discordant -- phred33 -I 10 -X 700.* Separate alignments were performed to spike-in *E. Coli* DNA (MG1655, Genbank U00096.3), from which sequencing depth was calculated and reads normalized accordingly before filtering into separate human and viral bedgraph files. Data quality control and visualization were performed using R. Data are available in the SRA database (PRJNA1047640).

Viral gene promoter coordinates previously identified for KOS strain (71) were BLAST sequenced against the 17Syn+ genome, and the equivalent region on the 17Syn+ genome used to generate viral promoter coordinates. Bedtools Mapbed was used to calculate the sum of scores at each defined promoter region, using normalized bedgraph files as input (promoter coordinates are listed in Table S3). Where a region lacked coverage, resulting in no score in the bedgraph file, a pseudovalue of 0.005 was used to allow fold enrichment calculation. Viral gene body coordinates were defined from the reference sequence NC_001806.2. Human promoters were located using the Eukaryotic Promoter Database (https://epd.expasy.org). The ratio between the sums for H3K27me2/3 and IgG was calculated to determine fold enrichment. MACS2 bdgcmp was used to generate linear fold enrichment bedgraph files for visualization, and Integrative Genome Viewer v2.16.1 was used to visualize bedgraph coverage files.

## Acknowledgments

This work was supported by The Owens Family Foundation (ARC), National Institutes of Health grants T32GM008136 (SAD and AKF), T32AI007046 (ALW), F32CA260116 (JH), MRC awards MC_UU_12014/5 and MC_UU_00034/2 (CB), and American Cancer Society award RSG-21-031-01-DMC (SBR). We thank the following for sharing plasmid constructs: Tobias Meyer (pLV-hTERT-IRES-hygro); Els Verhoeyen (pCMV-VSV-G, BaEVRLess); Philippe Mangeot (pBS-CMV-gagpol, BLADE); and Philippe Mangeot, Théophile Ohlmann, and Emiliano Ricci (BIC-Gag-Cas9). We thank David Knipe (Harvard Medical School) and Steven Triezenberg (Van Andel Institute) for sharing the RP5 and RP5R viruses and Sarah Dremel (National Cancer Institute) for advice and input on the CUT&RUN analysis.

---

<u>Figure S1 Histone peptide binding arrays for H3K27me3 and H3K27me2 antibodies show variable target specificity and non-specific binding affinities.</u>

Scatter plots of two binding array data sets from the same antibody sample, one dataset on each axis. Labels are bolded where the target residue is included. Other notable non-specific binding partners are also labeled.

<u>Figure S2 Validation of inhibitor activity and lack of interferon stimulated gene induction.</u>

(A) HFFs were treated with UNC1999 at indicated concentrations for 4 days, with fresh inhibitor added once on day 2. The cumulative effect of UNC1999 on cellular chromatin was assessed from histone extracts blotted for H3K27me3. Li-Cor band quantification is normalized to total H3 bands, relative to untreated cells. (B) Cumulative effects of treatment with GSK-J4 for four days on cellular chromatin, assessed by blotting histone extracts for both H3K27me3 and H3K9me3. Li-Cor band quantification in A and B was normalized to total H3 bands, relative to untreated cells. (C) IL-6 expression measured by RT-qPCR of cDNA made from HFFs treated with indicated concentrations of UNC1999 for 5 hours.

<u>Figure S3 The H3K27me2 antibody initially used in our experiments shows evidence of non-specific binding to the viral genome.</u>

(A) Comparison of co-localization with H3K27me2 immunostained with two different antibodies at 4 hpi (Kolmogorov-Smirnov test.) Adjusted p-value *=<0.05. (B) 17Syn+ genome coverage from HFFs 1 hpi, from a single replicate of CUT&RUN with control IgG and H3K27me2 antibodies (Diagenode C15410046-10).

