## Supplementary figures and images for "Single-genome analysis reveals heterogeneous association of the Herpes Simplex Virus genome with H3K27me2 and the reader PHF20L1 following infection of human fibroblasts"

### Supplemental Figure 1

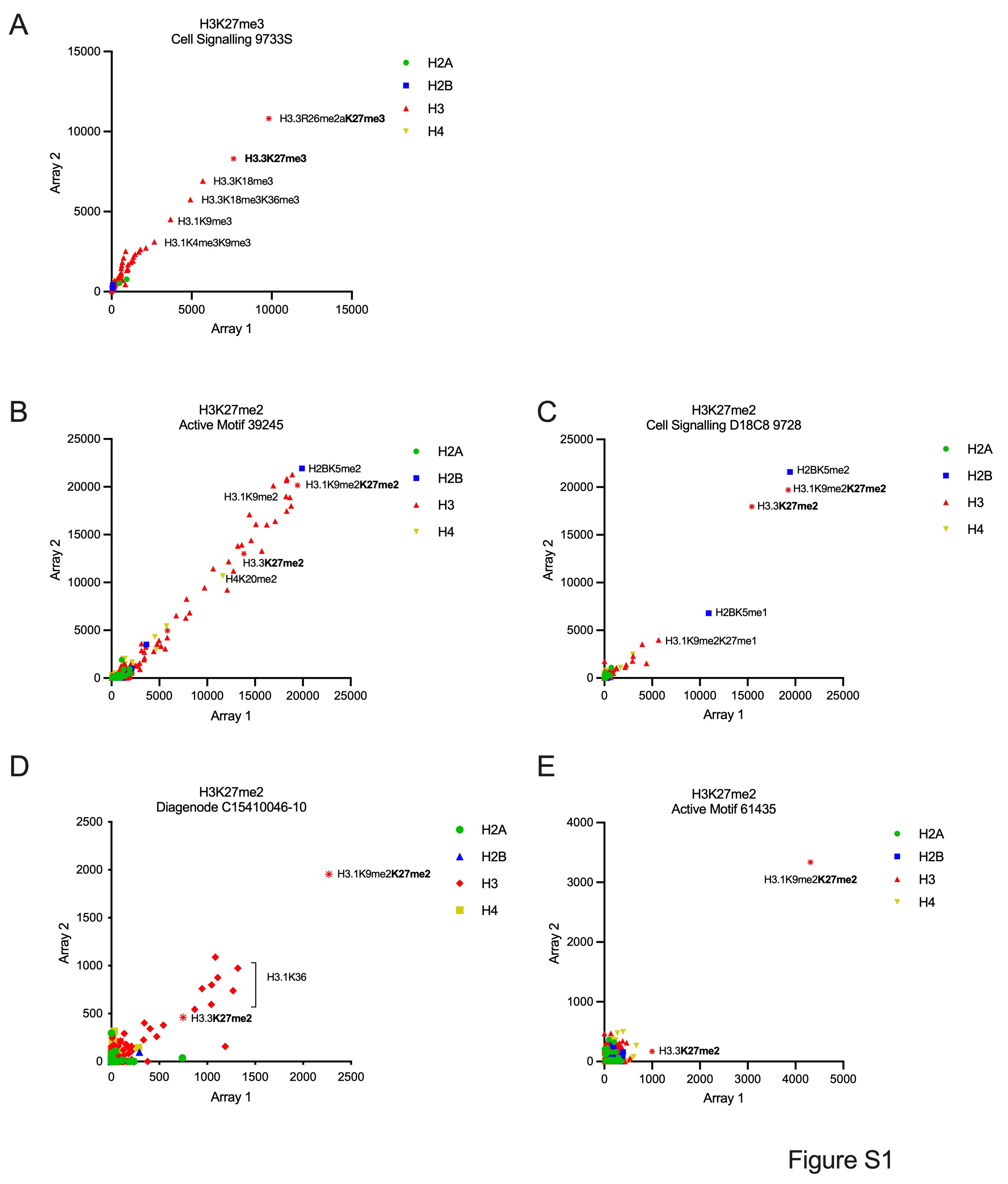

### Supplemental figure 2

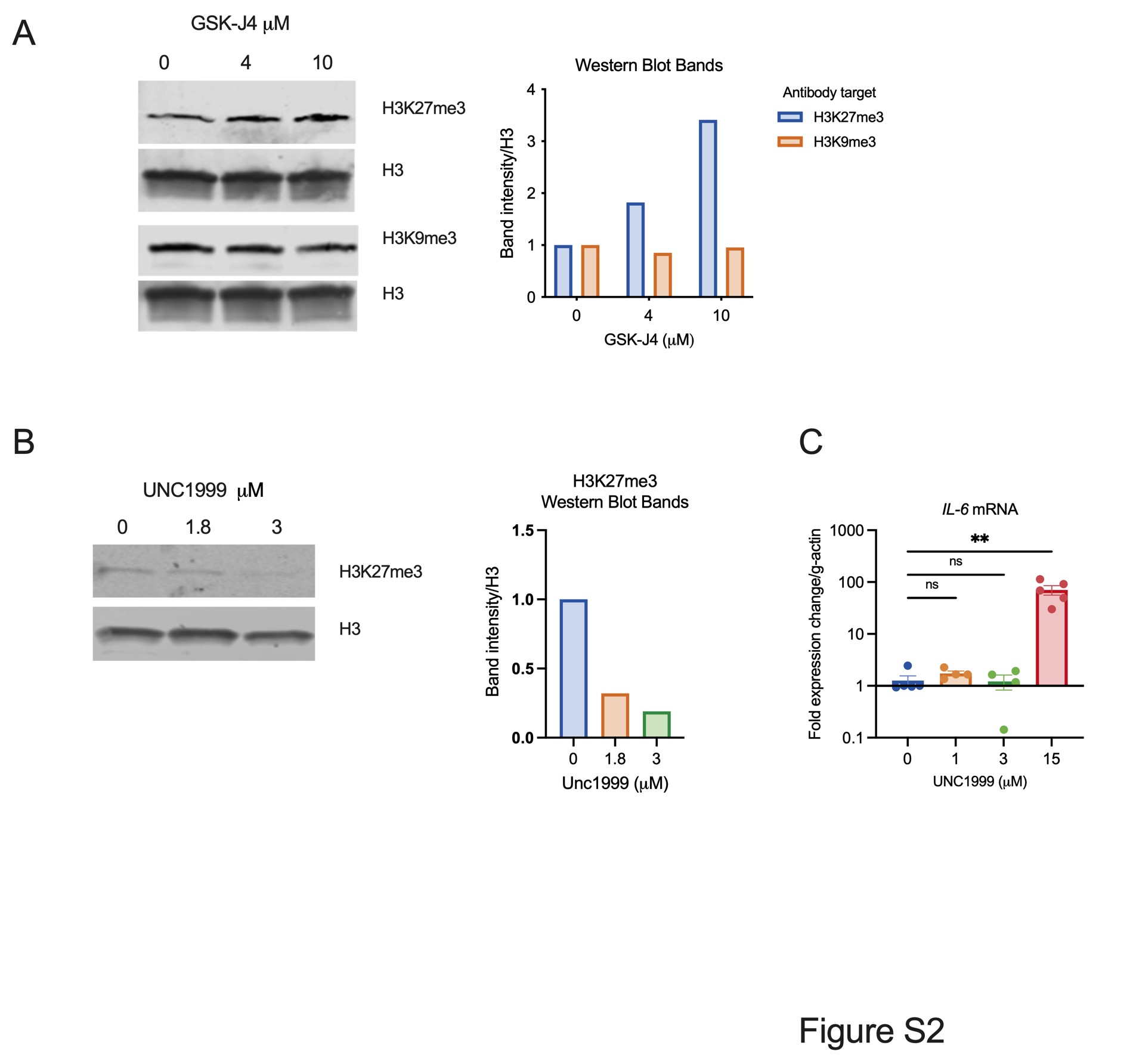

### Supplemental Figure 3

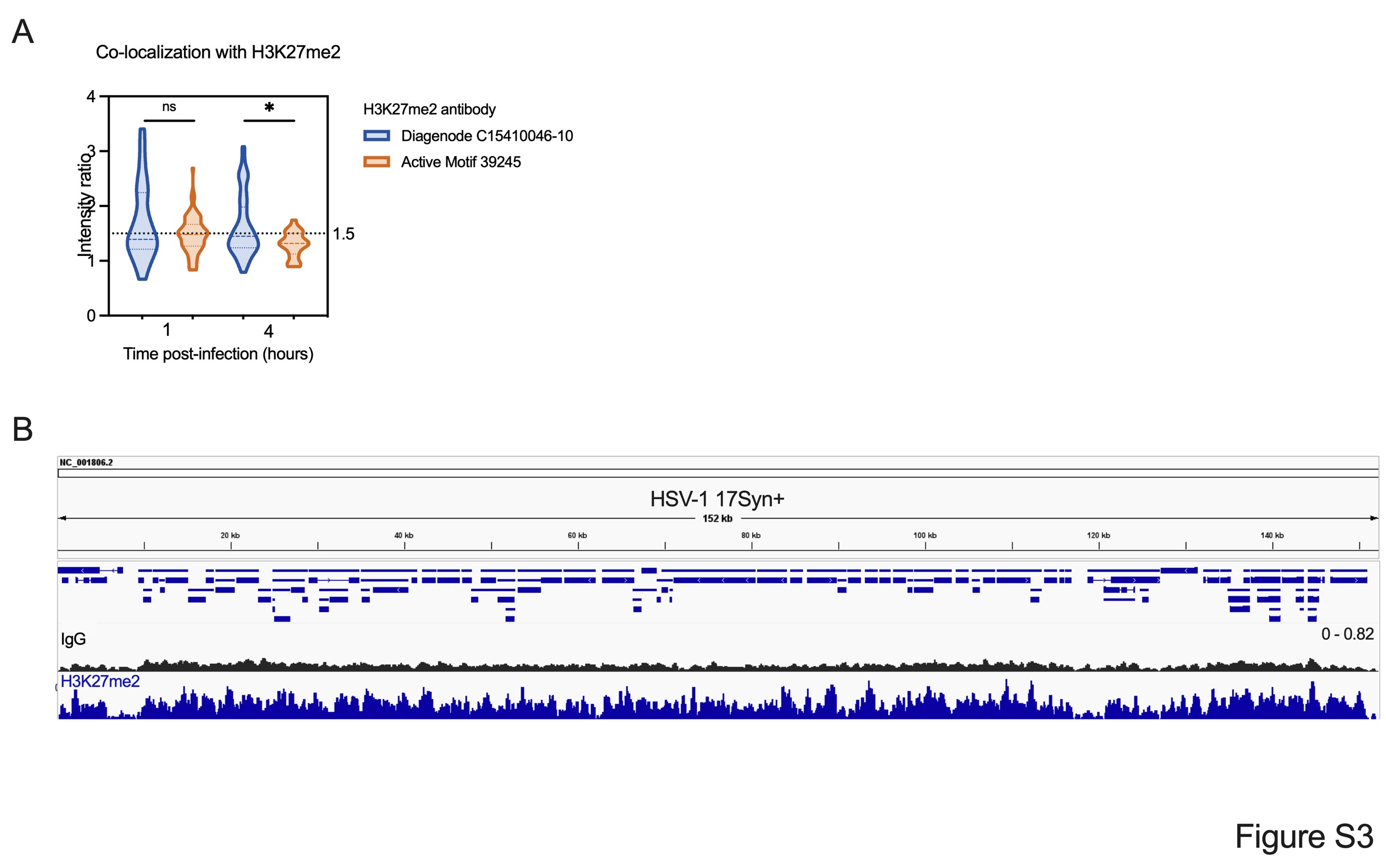
